# Phenotypic and transcriptomic analysis reveals early stress responses in transgenic rice expressing Arabidopsis DREB1a

**DOI:** 10.1101/2022.01.12.475916

**Authors:** Yasmin Vasques Berchembrock, Bhuvan Pathak, Chandan Maurya, Flávia Barbosa Silva Botelho, Vibha Srivastava

## Abstract

Overexpression of Arabidopsis Dehydration Response Element Binding 1a (*DREB1a*) is a well-known approach for developing salinity, cold and/or drought stress tolerance. However, understanding of the genetic mechanisms associated with *DREB1a* expression in rice is generally limited. In this study, *DREB1a* associated early responses were investigated in a transgenic rice line harboring cold-inducible *DREB1a* at a gene stacked locus. While the function of other genes in the stacked locus was not relevant to stress tolerance, this study demonstrates *DREB1a* can be colocalized with other genes for multigenic trait enhancement. As expected, the transgenic lines displayed improved tolerance to salinity stress and water withholding when compared to non-transgenic controls. RNA sequencing and transcriptome analysis showed upregulation of complex transcriptional networks and metabolic reprogramming as *DREB1a* expression led to the upregulation of multiple transcription factor gene families, suppression of photosynthesis and induction of secondary metabolism. In addition to the detection of previously described mechanisms such as production of protective molecules, potentially novel pathways were also revealed. These include jasmonate, auxin, and ethylene signaling, induction of *JAZ* and *WRKY* regulons, trehalose synthesis and polyamine catabolism. These genes regulate various stress responses and ensure timely attenuation of the stress signal. Furthermore, genes associated with heat stress response were downregulated in *DREB1a* overexpressing lines, suggesting antagonism between heat and dehydration stress pathways. In summary, through a complex transcriptional network, multiple stress signaling pathways are induced by DREB1a that presumably lead to early perception and rapid response towards stress tolerance as well as attenuation of the signal to prevent deleterious effects of the runoff response.

## INTRODUCTION

*Arabidopsis thaliana* transcription factor, Dehydration Response Element Binding 1a (DREB1a), is induced by cold, drought and salinity and considered highly promising for engineering abiotic stress tolerance in plants (Smirnoff and Bryant, 1999). Although it’s constitutive overexpression, is deleterious to plant growth, its conditional expression under abiotic stress inducible *Arabidopsis thaliana RD29a* promoter minimally affects plant growth and confers stress tolerance (Kasuga et al., 1999; Ito et al., 2006). Further, since *RD29a* is a stress-induced gene, it could promote *DREB1a* expression sustainably during stress conditions. *RD29a* promoter induced *DREB1a* transcripts to 2 – 10-fold higher under salinity, cold, and drought conditions in Arabidopsis, rice, and peanut (Latha et al., 2019; Oh et al., 2005; Sarkar et al., 2016), and transgenic expression of *RD29a:DREB1a* conferred tolerance to salinity, drought, and cold stress in diverse plant species including rice (Datta et al., 2012; Ganguly et al., 2020; Geda et al., 2019; Latha et al., 2019; Muthurajan et al., 2021; Oh et al., 2005; Ravikumar et al., 2014;), peanuts (Bhalani et al., 2019; Bhatnagar-Mathur et al., 2007; Devi et al., 2011; Sarkar et al., 2016) chickpea (Anbazhagan et al., 2015), soybean (de Paiva et al., 2014), chrysanthemum (Hong et al., 2006), potato (Behnam et al., 2007; Shimazaki et al., 2016) and *Salvia* sp. (Wei et al., 2016). It is now well accepted that enhanced expression of *DREB1a* by stress-regulated promoters such as *RD29a* is an effective strategy for conferring abiotic stress tolerance. However, whether a single gene will be sufficient to manage field level stress conditions is debatable. It is likely that stress tolerance genes such as *DREB1a* will have to be co-located with other genes that either confer stress tolerance or help overcome growth or yield trade-offs imposed by stress-response genes (Kudo et al., 2019; Shailani et al., 2021).

In the present study, transgenic rice developed through *Cre-lox* recombination mediated site-specific integration of multigene cassette were used (Srivastava & Ow, 2002). The rice lines contained *RD29a:DREB1a* gene along with other genes and the Cre-*lox* method facilitated precise full-length integration of the multigene cassette into a pre-determined genomic locus (Pathak & Srivastava, 2020). Thus, the functionality of *DREB1a* in japonica rice was tested from a stacked locus consisting of 4 other genes (marker genes).

Under the stress condition, Arabidopsis DREB1a interacts with *cis*-acting *DRE* (Dehydration Responsive Elements: RCCGAC; where R = A/G) for the regulation of stress responsive genes (Dubouzet et al., 2003; Yamaguchi-Shinozaki & Shinozaki, 1994). In Arabidopsis, its overexpression induces various stress responsive genes that contain *DRE* such as *rd17/cor47, erd7/10, erd10, cor15a/b cor15b, kin1, kin2/cor6.6, rd29A/cor78, Xero2* and *ADS2* in response to drought, salinity, and cold stress (Kasuga et al., 1999; Maruyama et al., 2004). However, in spite of its popularity in crop biotechnology, only a limited investigation has been done to reveal the downstream targets of DREB1a in other plant species such as rice. Microarray analysis of *japonica* rice seedlings expressing *ZmUbi1:DREB1a* (strong constitutive expression) found potential DREB1a targets but didn’t shed light on the mechanisms associated with drought/salinity tolerance (Oh et al., 2005). Ito et al. (2006) found proline and various sugar accumulation in rice overexpressing *DREB1a* or its rice ortholog (*OsDREB1a*) and proposed accumulation of these molecules as part of the stress tolerance mechanism. They also identified a few upregulated genes in the *DREB1a* transgenic rice. Finally, Latha et al. (2019) using microarray showed induction of *DRE*-containing genes encoding chloroplast structure and function in *RD29a:DREB1a indica* rice exposed to drought at reproductive stage. Similarly, RNA-seq analysis of *Salvia sp.* expressing *RD29a:DREB1a* showed induction of genes involved in photosynthesis, signal transduction and transcriptional activation, as well as carbohydrate transport, metabolism and protein protection (Wei et al., 2016). Finally, a study on *DREB1a* overexpressing chrysanthemum found similarities between DREB1a regulon of chrysanthemum and that of rice and Arabidopsis. This study used suppressive subtractive hybridization approach for identifying DREB1a targets in chrysanthemum (Ma et al., 2010).

Here, we analyzed transgenic rice lines in Taipei-309 background that contained a stack of genes, including *RD29a:DREB1a*. Taipei-309 is a chilling tolerant but drought and salinity sensitive *japonica* rice (Li et al., 2010). Accordingly, drought and salinity stress tolerance were studied at the young vegetative stages in the artificial media or greenhouse. The transgenic lines expressing *RD29a:DREB1a* showed significantly improved tolerance to both salinity (100 – 150 mM NaCl) and drought (10 – 20% soil water content) in comparison to the non-transgenic controls. Transcriptomic analysis of transgenic lines showed that induction of *DREB1a* led to the onset of complex transcriptional reprogramming along with hormone signaling and accumulation of a variety of protective molecules that protect the cellular components from the damaging effects of the stress. As some of these mechanisms have been reported in drought/cold/salinity stressed plants (Yamaguchi-Shinozaki & Shinozaki, 2006; Zhang et al., 2021), their enrichment in *DREB1a* overexpressing lines indicates involvement of multiple pathways and cross-talks among various transcription factors during early phase of the exposure to adverse condition towards helping the plant combat the stress.

## MATERIALS AND METHODS

### Plant materials

The previously described eight independent multigene stacked lines of rice cv. Taipei-309 were used in the present study for cold, drought and salinity stress. These lines contained *RD29a:DREB1a* gene in addition to 4 marker genes each expressed by strong constitutive or inducible promoters. These lines were developed using a multigene vector, pNS64, in the T5 founder line using Cre-*lox* mediated site-specific integration approach (Srivastava & Ow, 2002). Since each multigene stacked line is isogenic, as they harbored identical transgene integration and genomic location pattern, T1 seeds from homozygous lines carrying biallelic integration of pNS64 construct or the hemizygous lines containing monoallelic integration were pooled separately. Here, phenotypic data from homozygous lines is presented. For simplicity, the multigene transgenic lines harboring *RD29a:DREB1a* are referred to as ‘transgenic’ (T) and the parental T5 line as ‘non-transgenic’ (N) in this study.

### Salinity stress analysis

Two genotypes, *RD29a:DREB1a* transgenic line (T) and the non-transgenic (N) line as negative control, were submitted to salinity stress at different concentrations of sodium chloride (NaCl). Rice seeds were germinated on ½ MS medium solidified with 20% Phytagel (2g/L), and the seedlings at S3 stage, *i.e.*, having a prophyll emerged from coleoptile, were transferred to glass tubes containing ½ MS medium supplemented with 0 mM (control), 100 mM or 150 mM NaCl. The seedlings were kept under light intensity of 20 - 40 micromole/m^2^/s at 28°C until third-leaf stage (V3) (Moldenhauer et al., 2018). The experiment was conducted in complete randomized design in a factorial scheme (2×3), two genotypes in three salinity conditions, with fifteen replications and plot of a single plant. The number of chlorotic and necrotic leaves were evaluated daily, and the growth rate (GR) of the seedlings was evaluated every three days using the formula: GR = *l_x_* — *l_y_*, in which *l_x_* is the length (cm) on the day and *l_y_* is the length (cm) observed in the last measurement. The data was statistically analyzed by analysis of variance (ANOVA), and the Tukey test for comparisons among treatment means was applied with 5% as the level of significance using PROC MIXED of SAS (SAS, 2013) and R (R Core Team, 2017) software.

### Water withholding stress analysis

Seeds of the *RD29a:DREB1a* transgenic (T) lines and non-transgenic (N) controls were germinated in growth medium until the S3 stage when it was transferred individually to pots of 0.6 L containing 45g of a dry mixture of sphagnum peat moss and perlite (9:1), PRO-MIX LP15®. Before sowing, each pot was weighed, saturated with water, and placed in a tray filled with water for two days. The pots were then removed from the tray and placed on a grid until the water drained. When no dripping was observed, the pots were weighed again and this weight was considered as the field capacity (100% of soil water content, SWC). The weight of SWC at the field capacity was calculated as the difference between the pot weight with water-saturated soil and with dry soil (Almeida et al., 2016). All the plants were kept at field capacity for two weeks, when half of the plants were subjected to water stress. The irrigation in these plants was suspended and the pots were weighed every day until they reached 10% of SWC (representing a severe stress). Subsequently, the plants were irrigated again for one week for recovery. The plants were kept in growth chamber at 28°C and 300–500 μE/m 2/s light intensity with a 12-h photoperiod during all the experiment. A completely randomized design in a factorial scheme (2×2) with ten replications was used with each plot consisting of one plant. The two treatments consisted of two water regimes, well-watered control (100% SWC) and water stressed condition (10% SWC). The plants in stress were checked every day to determine the SWC at which the plants started to display the first symptoms of withering, chlorosis, and leaf senescence. After one week of recovery, all the plants were analyzed for percentage of leaves with chlorosis and percentage of leaves with necrosis. The phenotypic data was statistically analyzed by analysis of variance (ANOVA), and the Tukey test for comparisons among treatment means was applied with 5% as the level of significance using PROC MIXED of SAS (SAS,2013) and R (R Core Team 2017) software.

### RNA-seq

Seeds of the transgenic (T) and non-transgenic controls (N) were germinated on ½ MS media and the 10 days old seedlings were divided into two groups: cold-shock treatment (CS) and the control treatment at the room temperature (RT). Six CS-or RT-treated seedlings (biological replicates) were individually used for total RNA isolation using Trizol reagent (Thermo Fisher Scientific, USA). RNA was treated with DNase I and its concentration and quality were measured with Qubit 2.0 fluorometer (Thermo Fisher Scientific, USA). Subsequently, RNA samples of all 6 replicates were pooled for each genotype and treatment yielding four pooled groups: T_CS, T_RT, N_CS, and N_RT. cDNA library synthesis and Illumina sequencing was done at Novogene Inc. RNA quality and quantity was determined by High Sensitivity RNA TapeStation (Agilent Technologies Inc., California, USA) and Qubit 2.0 RNA High Sensitivity assay (Thermo Fisher Scientific, USA), respectively. Paramagnetic beads coupled with oligo d(T)25 were combined with total RNA to isolate poly(A)+ transcripts based on NEBNext^®^ Poly(A) mRNA Magnetic Isolation Module manual (New England BioLabs Inc., Massachusetts, USA). All libraries were constructed according to the NEBNext® Ultra™ II Directional RNA Library Prep Kit for Illumina^®^ (New England BioLabs Inc., Massachusetts, USA). The libraries were quantified by Qubit 2.0 and their quality was assessed by TapeStation D1000 ScreenTape (Agilent Technologies Inc., California, USA). Final library average size was about 390 bp with an insert size of about 270 bp. Illumina^®^ 8-nt dual-indices were used for paired end sequencing. Equimolar pooling of libraries was performed based on QC values and sequenced on an Illumina^®^ HiSeq (Illumina, California, USA) with a read length configuration of 150 paired-end for 40 million paired-end reads per sample (20 million in each direction). The data discussed in this publication have been deposited in NCBI’s Gene Expression Omnibus (Edgar et al., 2002) and are accessible through GEO Series accession number GSE185088 (https://www.ncbi.nlm.nih.gov/geo/query/acc.cgi?acc=GSE185088).”

### Differential Gene Expression Analysis

The differential gene expression (DEG) analysis was performed using main GALAXY platform (usegalaxy.org) version 2.7.5b (Afgan et al., 2018). The raw data quality in the fastq format was assessed using FastQC tool. Further quality check and removal of adapter sequences were done by TRIMMOMATIC tool using SLIDINGWINDOW, MINLEN, and AVGQUAL parameters. The raw reads were mapped on RNA-STAR using rice genome (*Oryza sativa* japonica Group) downloaded from Ensemble version 48 in the FASTA format (ftp://ftp.ensemblgenomes.org/pub/plants/release48/fasta/oryza_sativa/dna/Oryza_sativa.IRGSP-1.0.dna.toplevel.fa.gz). The Ensemble version 48 cDNA annotation in GTF format (ftp://ftp.ensemblgenomes.org/pub/plants/release-49/gtf/oryza_sativa/Oryza_sativa.IRGSP-1.0.48.gtf.gz) were used for generating the genome indexing. The normalized counts were obtained from Featurecounts version 1.6.4 (Liao et al., 2014) using Gene-ID”t” read-count, gene_id features. The differential gene expressions (DEG) were obtained using edgeR with log2 fold change ≥ 2, P-value ≤0.01, and P-value adjustment method of Benjamin and Hochberg (Liu et al., 2015). The genes were considered differentially expressed if values were above the threshold (log2 fold change ≥ 2 or ≤-1 and FDR ≤0.01) in the comparison of cold-stressed *RD29a:DREB1a* transgenic lines (T_CS) versus cold-stressed non-transgenic control (N_CS) and in roomtemperature *RD29a:DREB1a* transgenic lines (T_RT) versus room-temperature non-transgenic controls (N_RT). To determine the RNA read counts for transgenes stacked in the transgenic line, the aligned reads from T_CS and T_RT were filtered using BAM Split tool into mapped and unmapped reads. The unmapped reads were mapped against *NPTII* (U55762.1), *GFP* (U55762.1), *GUS* (AF485783), *DREB1a* (NM_118680.2), and *pporRFP* (DQ206380.1) sequences downloaded from NCBI in the fasta format using “map with BWA version” 0.07.17.4 (Li & Durbin, 2010) containing “IS” algorithm mode. These aligned reads were sorted using Samtools, and reads were obtained using Samtools idxstat. Heatmaps of DEGs, k-means clusters, and gene network were developed using iDEP.93 (http://bioinformatics.sdstate.edu/idep93/; Ge et al., 2018). Gene Ontology (GO) for the upregulated and downregulated DEGs was performed using BLAST2GO against nucleotide (NR) database (Conesa et al., 2005; Conesa & Gotz, 2008) and gene set enrichment analysis (GSEA) was carried out using gProfiler version Ensembl 103 (Reimand et al., 2016) (https://biit.cs.ut.ee/gprofiler/gost) using hypergeometric distribution adjusted by set count size (SCS) for multiple hypothesis correction for removal of enriched false positive GO terms. KEGG analysis was performed for the downregulated and upregulated genes in BLASTKOALA (https://www.kegg.jp/blastkoala/; Kanehisa et al., 2016).

### Search of DRE elements

Drought response element (DRE) in DEGs were searched using PLANT PAN 3.0 (Support ≥ 70% frequency of promoters containing the TF; Chow et al., 2019). For this purpose, 2 kb sequence upstream of the start codon of 683 up-regulated genes (log2 fold change ≥ 2 or ≤-1 and FDR ≤0.01) were obtained from BioMart in Plant Ensemble v48. The extracted regions of these genes were searched for DRE core-motif (A/GCCGAC) #292.

### qRT-PCR

The pooled RNA for each genotype and treatment was used for qRT-PCR. One microgram DNAse-treated RNA was converted to cDNA using Prime RT-PCR kit (Takara Bio Inc.). The qRT-PCR was performed on 11 selected DEGs containing DRE elements. The primers for each gene are listed in **Supplementary Table 1** with *Os7Ubiquitin* serving as the reference gene. All reactions were performed in two technical replicates and fold-change was calculated by 2^ddCT method (Livak & Schmittgen, 2001).

## RESULTS

### Seedling response to salinity stress

*RD29a:DREB1a* transgenic (T) lines, and non-transgenic (N) controls were tested for the salt stress assay with two different concentrations of NaCl. Significant genotype effect (p<0.05) was observed for all evaluated traits, indicating a difference between T and N lines. At the beginning of vegetative stage (3^rd^ day), the T seedlings showed the same growth rate under non-stress (0mM NaCl) or stress (100mM NaCl) conditions (**Figure 1a**). Although T lines grew faster than the N controls under 0 and 100 mM NaCl, a similar growth rate was observed for both genotypes in 150mM NaCl, indicating a severe effect of higher salinity condition. The N seedlings had a negative response to stress, decreasing 20.2% of growth rate when compared salt control (0mM) and 100mM NaCl. The maximum reduction in growth rate in stressed seedlings was observed on 6^th^ day for both T and N lines (**Figure 1b**). In 100mM NaCl, there was a growth rate reduction of 4-fold and 19-fold, respectively, for T and N in comparison to salt control (0mM NaCl). However, no significant difference was found in growth rate of T and N seedlings under more severe stress (150mM NaCl). On the 9^th^ day, the growth rate for T plants was greater than N plants for both salinity stress conditions (**Figure 1c**). Although T seedlings had a small reduction in growth rate under 100mM, their performance was similar to N seedlings under no stress (0mM). The N lines, in turn, were more sensitive to salinity showing a reduction of 2.9-fold under 100 mM, and 3.6-fold under 150mM. On the 12^th^ day, growth rate slowed down (**Figure 1d**), however, performance of T plants under salt stress was similar to those under control (0 mM NaCl), whereas N plants showed a significant reduction in growth in 150mM NaCl.

**Figure 1:**
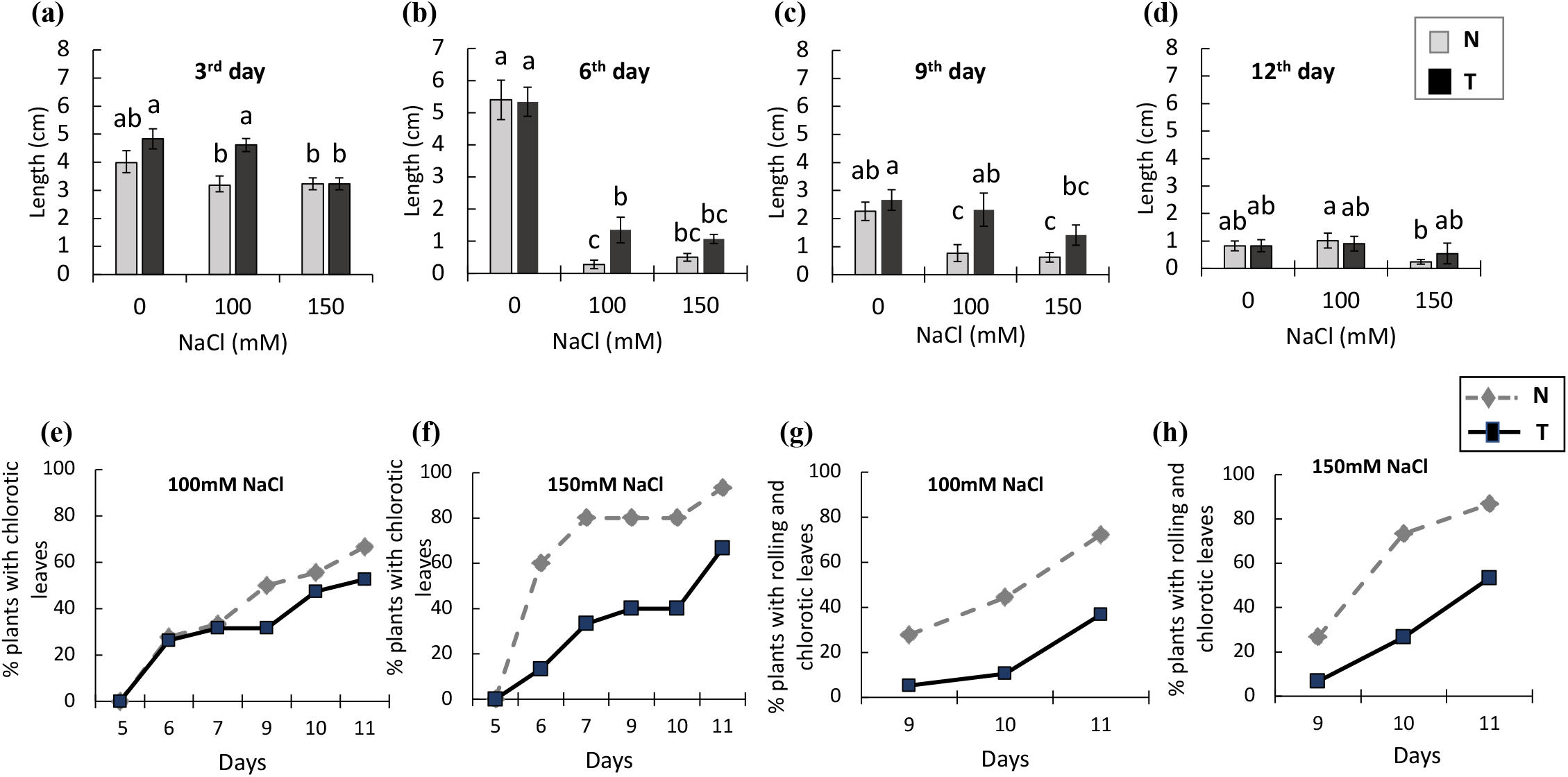
Salinity stress analysis on *RD29a:DREB1A* transgenic lines (T) and non-transgenic (N) controls under 0mM, 100mM and 150mM NaCl concentrations. (**a-d**) Seedling growth rates recorded on 3rd (a), 6th (b), 9th (c), and 12th day (d) of treatment. (**e-f**) Percentage of plants with chlorotic leaves under salinity stress of 100mM (e) and 150mM NaCl (f); (**g-h**) Percentage of plants with rolling and chlorotic leaves under salinity stress of 100mM (g) and 150mM NaCl (h). Different letters indicate differences by Tukey test (n=15). The error bars indicate standard errors. N and T genotypes are shown as gray or black bars, respectively, in (a-d) and dashed and solid lines in (e-h).

Leaf chlorosis started to be observed from 5^th^ day of stress, for both 100mM and 150mM NaCl (**Figure 1e-f**). Both the genotypes had 30% of seedlings with chlorotic leaves on the seventh day in 100mM NaCl. However, this percentage increased more sharply over the days in N lines, showing, at the end, 14% more plants with chlorosis than T lines (**Figure 1e**). In 150mM NaCl, this difference was greater and could be observed since the 6^th^ day, reaching in 93% and 67% in N and T seedlings, respectively (**Figure 1f**). The percentage of chlorotic leaves that also showed rolling, a symptom of severe stress, was greater in N plants with a difference of 35% (100mM NaCl) and 34% (150mM NaCl) compared to T plants (**Figure 2g-h**).

**Figure 2:**
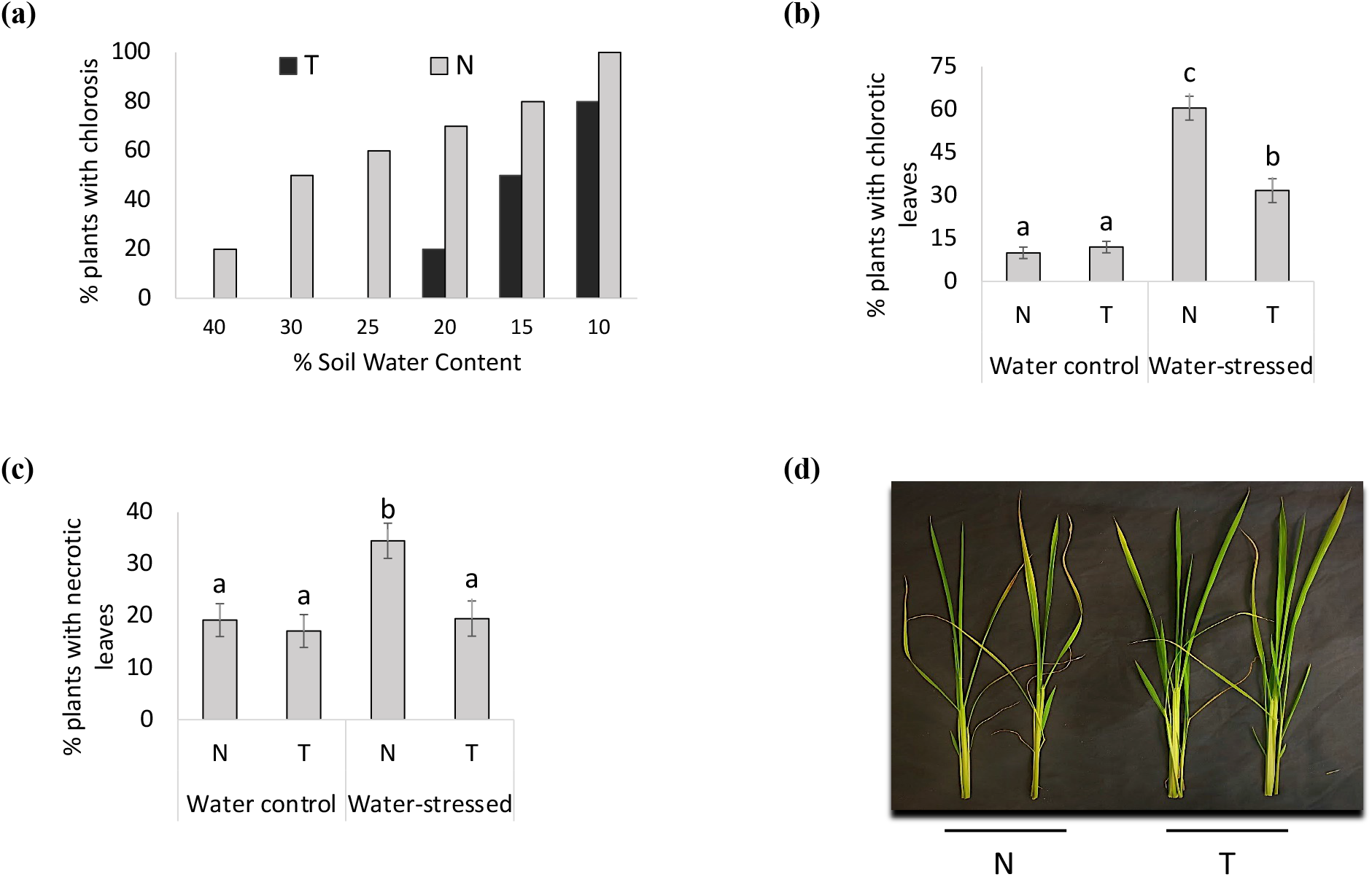
Water withholding stress analysis of *RD29a:DREB1A* transgenic lines (T) and non-transgenic (N) controls. (**a**) Percentage of plants with chlorotic leaves under different soil water contente (SWC), (**b-c**) Percentage of plants with chlorotic (b) and necrotic (c) leaves at the end of the experiment in water control (100% SWC) and water stressed (10% SWC) plants, and (**d**) Shoots of N and T plants after 7 days of water recovery. Different letters indicate differences by Tukey test (n=10). The error bars indicate standard errors.

### Plants response to drought stress

After submitting the rice plants to drought stress, the phenotypic symptoms caused by stress were evaluated in the vegetative stage. After withholding water, the plants from N line started to show the first symptoms of stress in leaves (chlorosis) at 40% of soil water content (SWC), while no symptoms were observed in T plants (**Figure 2a**). At 20% SWC, 70% of N plants exhibited the leaf chlorosis as the symptom of stress, while T lines had just started to develop first symptoms of stress. The *DREB1A* gene seemed to have increased the tolerance to drought as majority of T plants were asymptomatic in 20% SWC. Further, in maximum stress (10% SWC), a few T plants looked normal without any apparent symptoms, while all N plants showed chlorosis (**Figure 2a**). At the end of the experiment (at 10% SWC), plants showing chlorotic and necrotic leaves were 1.9-fold and 1.8-fold lower, respectively, in T lines as compared to N line (**Figure 2b-c**). Finally, N plants had more wilted and chlorotic leaves even after the water recovery time than T plants (**Figure 2d**).

### Transcriptome analysis of cold-stress induced DREB1a transgenic lines

To understand the potential mechanism by which DREB1a provides stress tolerance to rice, RNA-seq was done on cold-stressed (CS) and room temperature controls (RT) of two genotypes: transgenic line expressing *RD29a:DREB1a* (T) and non-transgenic controls (N) (**Table S2**). The processed reads were mapped to the rice genome and the differentially expressed genes (DEGs) were identified through comparisons between genotypes (T *vs* N) in the respective treatments (CS *vs* RT). *RD29a* promoter is induced by multiple abiotic stresses including cold stress (Msanne et al., 2011). Earlier, transgenic rice lines #9, #11, and #29 were characterized to strongly induce *DREB1a* upon cold treatment (Pathak & Srivastava, 2020). Consistent with that, RNA-seq comparison of cold-stressed transgenic lines with the room temperature control (T_CS *vs* T_RT) showed ~49 fold higher read counts of *DREB1a*, while no significant change (<2 fold) was observed in other transgenes (marker genes) stacked in the same locus. (**Table S3**).

A comparison of T_CS *vs* N_CS and T_RT *vs* N_RT showed 2802 unique DEGs in T_CS, 109 unique DEGs in T_RT, and 121 DEGs common to T_CS and T_RT. (**Figure 3a**). The k-means clustering of top 2000 most-variable genes using normalized expression counts showed that T_CS contained a large cluster of strongly upregulated genes and a smaller cluster of down-regulated genes, while the two genotypes at room temperature (T_RT and N_RT) showed a similar expression pattern (**Figure 3b**). These observations fit the hypothesis that upon induction of *RD29a:DREB1a* by cold-shock (CS) several genes are induced through binding of DREB1a to the promoters of the target genes, and the leaky expression of *RD29a* promoter at RT results in the induction of a smaller set of the DREB1a targets at RT. The list of DEGs were further filtered with the criteria of p-value ≤0.01 and log2 fold change ≥2. The resulting 2069 genes were used for the gene ontology (GO) and pathway analysis.

**Figure 3:**
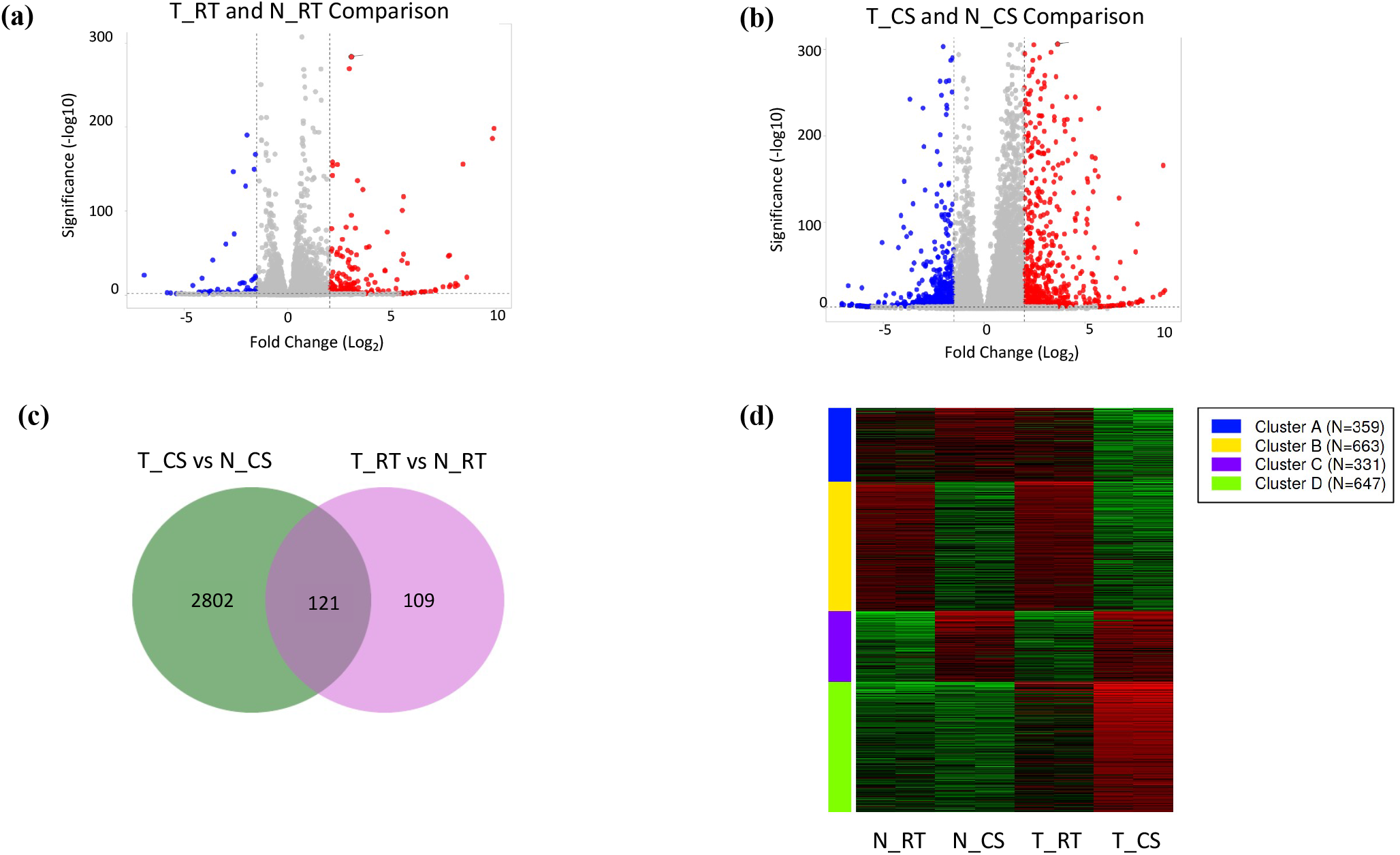
Differentially expressed genes (DEGs) in *RD29a:DREB1A* transgenic lines (T) upon 20 h cold-shock (CS) treatment (T_CS) or under room temperature (RT) control condition (T_RT) in comparison to non-transgenic (N) controls in respective treatments (N_CS or N_RT). (**a-b**) Volcano plot showing DEGs between two genotypes upon CS or RT treatments. Blue and red represent downregulated and upregulated genes, respectively, and dashed lines show log2 cut off for differential expression; (**c**) Venn diagram showing distribution of DEGs in T_CS and T_RT in comparison to N_CS and N_RT, respectively (Total number of genes= 31,468; FDR cut off 0.01, log2FC ≥2); (**d**) k-means clustering of top 2000 most-variable genes using normalized expression counts. Heatmap shows lower (green) to higher (red) expression of the cluster of genes in T_CS, N_CS, T_RT, and N_RT including two replicates of each genotype_treatment. Data analyzed on iDEP.93 http://bioinformatics.sdstate.edu/idep93/ (Ge et al., 2018).

### Functional annotation of genes induced in *RD29a:DREB1a* transgenic line upon cold stress

The GO assessment of 2069 DEGs expressed in T_CS relative to N_CS and 289 DEGs in T_RT relative to N_RT was performed using BLAST2GO. This analysis in T_CS corresponded to 836 GO terms in biological process (BP), 482 in molecular function (MF) and 171 in cellular components (CC), while in T_RT, 171, 161 and 41 GO terms were associated, respectively, with BP, MF and CC. The upregulated BP in T_CS included protein phosphorylation, signal transduction, response to chemical and hormone stimulus, and jasmonic acid mediated signaling, MF included kinase activity and protein Ser/Thr kinase activity, and CC included cell periphery and plasma membrane (**Figure 4a, b; Figure S1a-c**). The down regulated genes in T_CS, on the other hand, were associated with response to heat, protein folding, cell cycle process, and photosynthesis, light harvesting in photosystem I (**Figure S1a-c**). The gene set enrichment analysis (GSEA) showed that upregulated DEGs in T_CS were involved in protein serine/threonine kinase activity, kinase activity, catalytic activity, protein kinase activity, oxidoreductase activity and DNA binding transcription factor activity. The down-regulated genes in T_CS were involved in unfolded protein binding, microtubule binding, tubulin binding, protein heterodimerization activity, and microtubule motor activity. Among the biological pathways, plant hormone signal transduction and α-linolenic acid metabolism were highly enriched in the upregulated genes, and diterpenoid biosynthesis, DNA replication, photosynthesis-antennae proteins, and protein processing in endoplasmic reticulum pathways were enriched in downregulated genes. In T_RT, the upregulated genes were enriched in DNA-binding transcription factor activity, and transcription related processes, although, no pathways were significantly expressed. In the downregulated genes, no significant pathways were detected, however, two genes involved in cutin, suberine and wax biosynthesis were enriched (**Table S4**)

**Figure 4:**
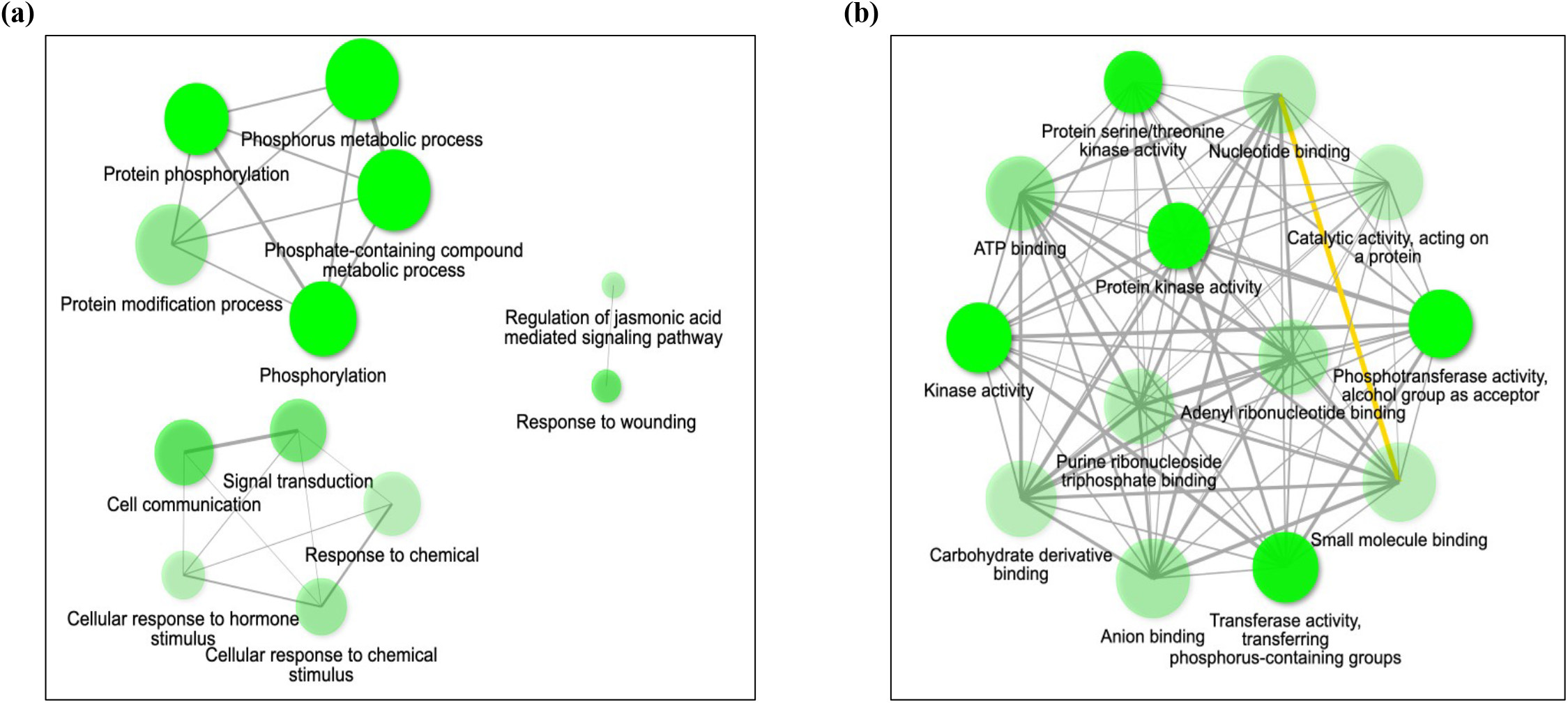
Network of upregulated pathways in *RD29a:DREB1a* transgenic line upon cold-shock in comparison to cold-shocked non-transgenic controls. (**a**) network of biological processes, and (**b**) network of molecular functions (FDR<0.01; log2FC ≥2). Data analyzed on iDEP.93 http://bioinformatics.sdstate.edu/idep93/ (Ge et al., 2018).

The Kyoto Encyclopedia of Genes and Genomes (KEGG) pathways were analyzed only for T_CS. This analysis showed that a higher percentage of down-regulated genes were associated with genetic information processing, whereas a higher percentage of upregulated genes were associated with carbohydrate metabolism, environmental information processing, lipid metabolism, signaling, and secondary metabolite synthesis among others (**Figure 5a**).

**Figure 5:**
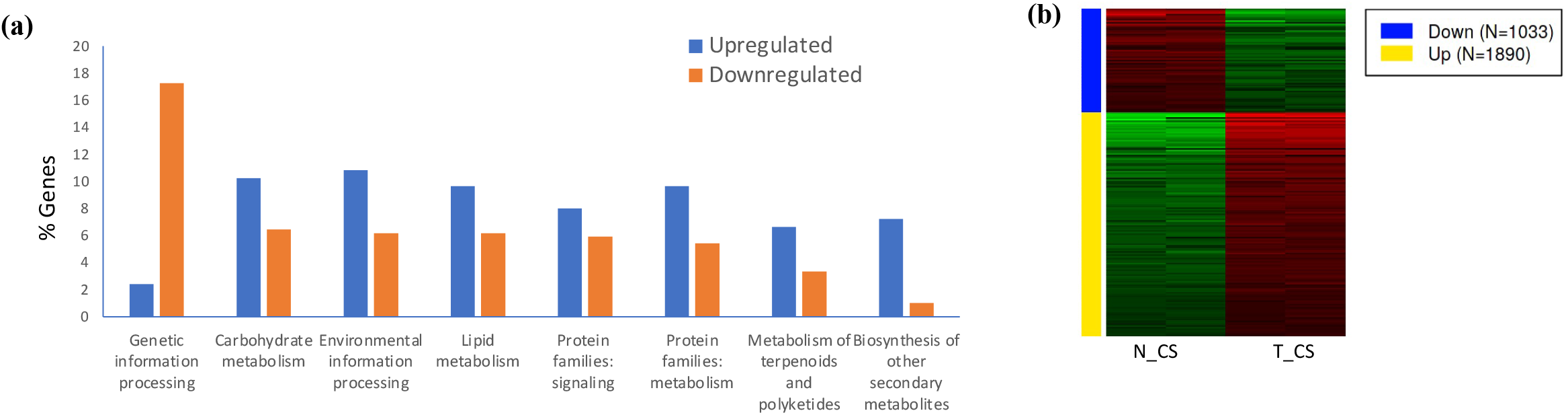
Rice genetic pathways regulated by DREB1a. (**a**) upregulated or downregulated Kyoto Encyclopedia of Genes and Genomes (KEGG) pathways in cold-shocked transgenic line expressing *RD29a:DREB1a* in comparison to cold-shocked non-transgenic controls. (**b**) Cluster of upregulated and downregulated genes in two technical replicates of transgenic (T) or non-transgenic (N) lines treated with cold-shock (CS) (FDR<0.01; log2FC ≥2). The list of upregulated genes was used for functional annotation and their association with pathways.

Corroborating with the GSEA, the upregulated genes were associated with phenylpropanoid biosynthesis, flavonoid biosynthesis, α-linolenic acid metabolism, and amino acids metabolism, while MAPK signaling, hormone signaling, and calcium signaling were among the upregulated environmental information processing pathways. In summary, GSEA and KEGG pathway analysis indicate induction of transcription factor (TF) genes that potentially regulate secondary metabolites and sugars metabolism in providing protective functions, stress-sensing, and hormone and calcium signaling towards preparing the plant against stress-induced damage.

The mechanisms by which DREB1a confers stress tolerance in rice was further investigated by analyzing the cluster of DEGs based on K-means clustering (FDR <0.01, log2FC >2) and the function of the top DEGs in T_CS vs N_CS comparison (**Figure 5b**). The two approaches converged to mostly common outcomes that shed light on likely pathways involved in the process (**Figure 6**). The most significant pathways are described below.

**Figure 6:**
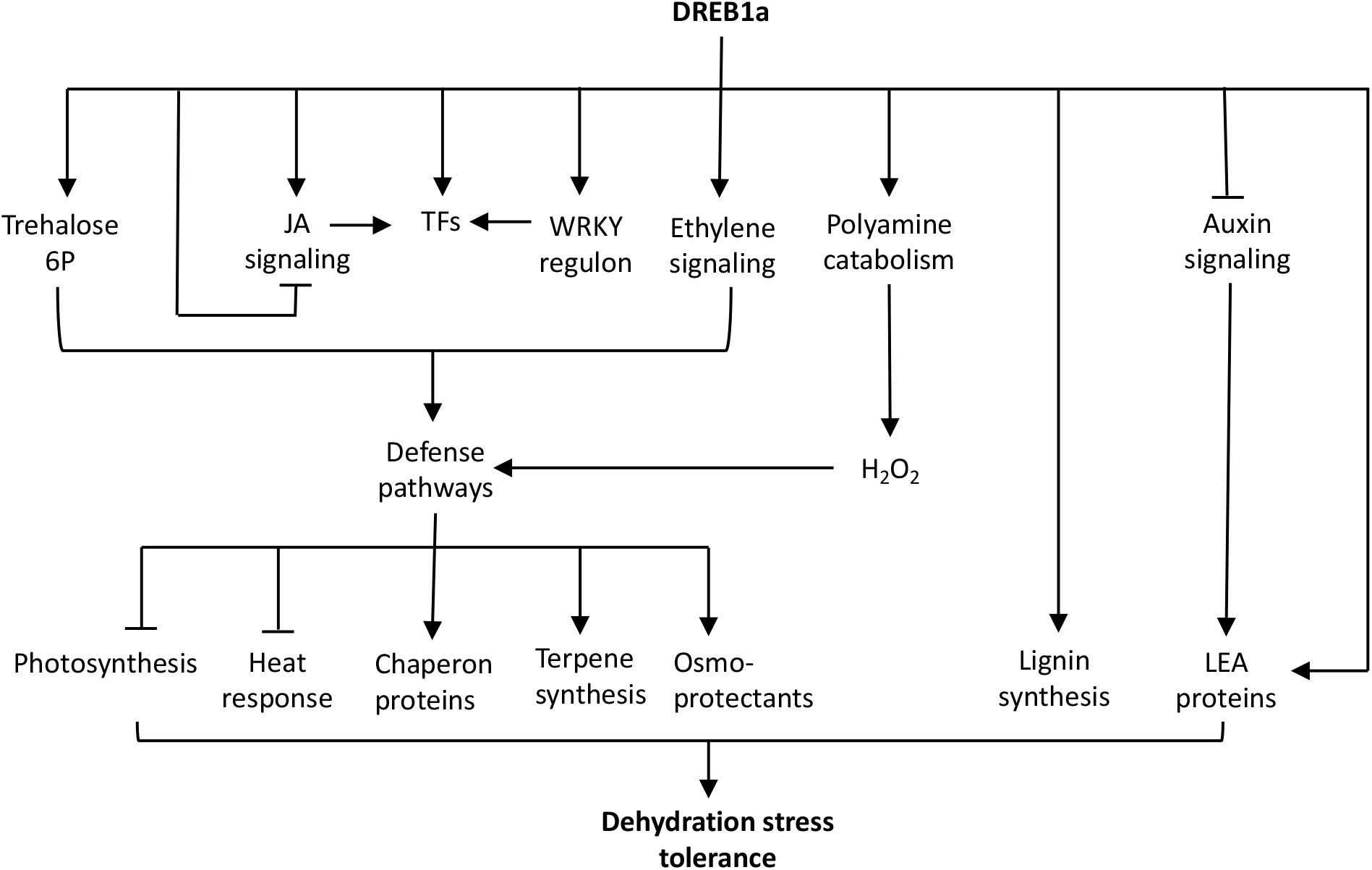
Proposed mechanism of dehydration stress tolerance in the rice seedlings expressing DREB1a. Sustained induction of DREB1a leads to the expression of transcription factor (TF) genes that positively regulate defense pathways. Production of signal molecules such as Trehalose-6-P and hydrogen peroxide (H_2_O_2_) through induction of genes encoding metabolic enzymes reinforce defense pathways. Through direct or indirect interactions, DREB1a induces lignin biosynthesis, production of osmo-protectants, secondary metabolites, and chaperon proteins that fortify cell wall, scavenge oxidants, and protect cellular membranes. To support stress response, resources are diverted from primary metabolism through suppression of photosynthesis and antagonistic pathways such as heat stress response, eventually leading to stress tolerance. Upregulations and suppressions of pathway/component is shown by arrows and stubbed lines, respectively.

### Jasmonic acid (JA) signaling

The GO analysis of the cluster of DEGs uniquely upregulated in T_CS indicated a role for JA mediated signaling and genes involved in response to wounding among others (**Table 1**). JA signaling is an important defense response in plants against insects, pathogens, and abiotic stresses (Wasternack, 2007). JAZ (JASMONATE ZIM-DOMAIN) proteins repress JA signaling but are subject to proteasome-mediated degradation by JA pulse treatment (Chini et al., 2007; Thines et al., 2007). Thus, degradation of JAZ is a key step in unleashing JA signaling or wound response. The top JA signaling genes in T_CS are all JAZ family of genes: *OsJAZ1, OsJAZ2, OsJAZ5, OsJAZ8/9*, and *OsJAZ11* (**Table 2**). However, since scant information is available on JAZ targets, understanding how *JAZ* induction leads to abiotic stress tolerance is not clear. It is possible that JAZ actively repress factors that negatively control stress response and the induction of *JAZ* is important in attenuating the response upon transmission of the signal. Of the 5 upregulated *JAZ*, *OsJAZ5* and *OsJAZ7* contained 3 *DRE* elements each. On the other hand, genes associated with α-linolenic acid metabolism, including lipoxygenases *(LOX1)* and 12-oxophytodienoic acid reductase *(OPR)* that participate in JA synthesis, are also upregulated in T_CS (**Table 2**). Some of *LOX1* genes and *OPR1* contain *DRE* elements. In summary, genes encoding JAZ and JA biosynthesis could serve as direct targets of DREB1a that regulate JA signaling as part of the stress response.

**Table 1:**
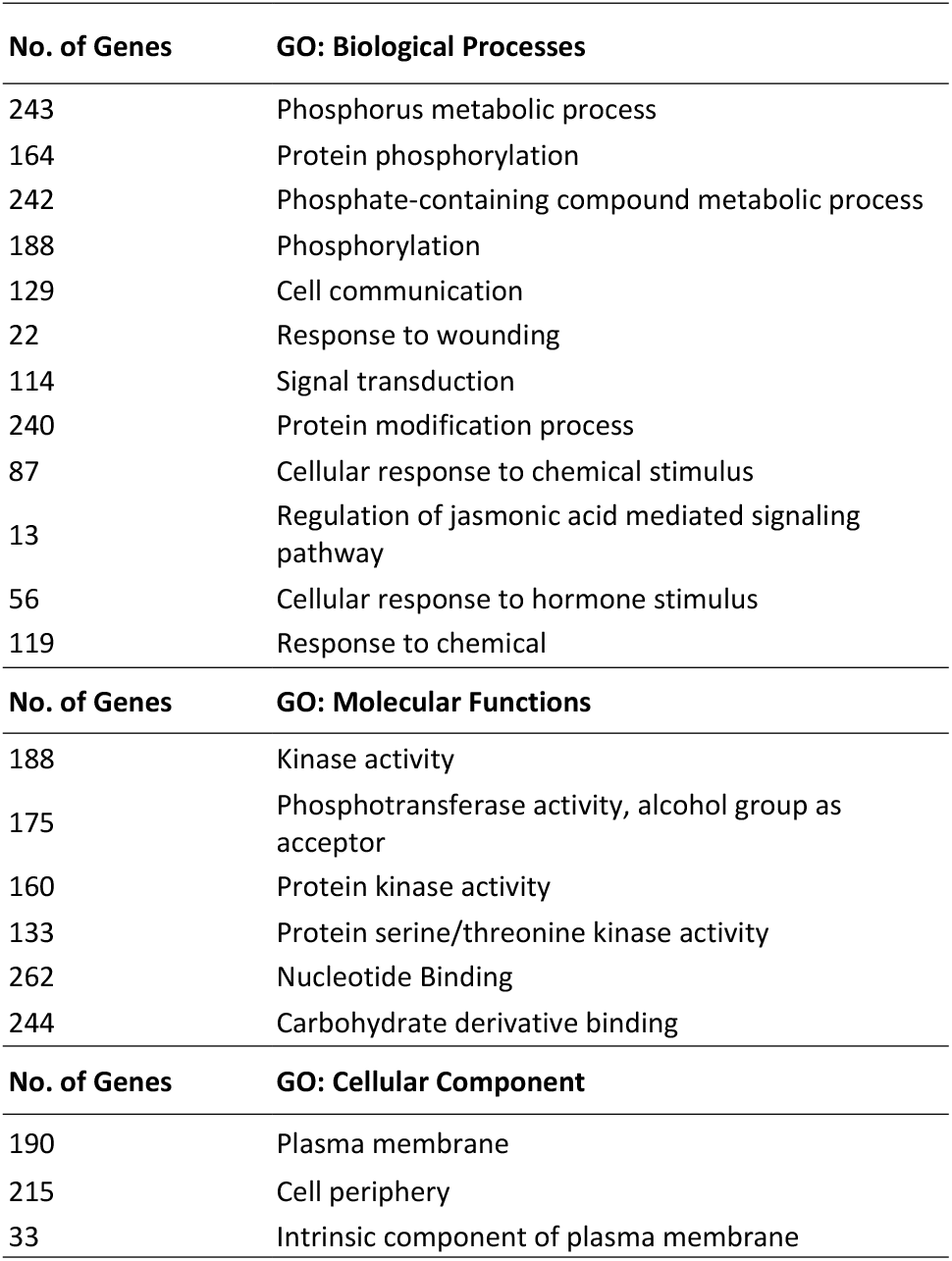
GO terms associated with upregulated genes in cold-stressed *RD29a:DREB1a* transgenic line (T_CS) relative to the cold-stressed non-transgenic line (N_CS)

| No. of Genes | GO: Biological Processes |
| --- | --- |
| 243 | Phosphorus metabolic process |
| 164 | Protein phosphorylation |
| 242 | Phosphate-containing compound metabolic process |
| 188 | Phosphorylation |
| 129 | Cell communication |
| 22 | Response to wounding |
| 114 | Signal transduction |
| 240 | Protein modification process |
| 87 | Cellular response to chemical stimulus |
| 13 | Regulation of jasmonic acid mediated signaling pathway |
| 56 | Cellular response to hormone stimulus |
| 119 | Response to chemical |
| No. of Genes | GO: Molecular Functions |
| 188 | Kinase activity |
| 175 | Phosphotransferase activity, alcohol group as acceptor |
| 160 | Protein kinase activity |
| 133 | Protein serine/threonine kinase activity |
| 262 | Nucleotide Binding |
| 244 | Carbohydrate derivative binding |
| No. of Genes | GO: Cellular Component |
| 190 | Plasma membrane |
| 215 | Cell periphery |
| 33 | Intrinsic component of plasma membrane |

**Table 2:** Biological pathways induced by DREB1a in rice

| Pathway/Function | Genes | Gene ID | Fold change | No. of DRE |
| --- | --- | --- | --- | --- |
| Jasmonic Acid signaling (JAZ regulon) | OsJAZ1/TIFY11d | Os10g0392400 | 3.6 | - |
|  | OsJAZ11/TIFY9 | Os04g0395800 | 5.4 | 3 |
|  | OsJAZ5/ TIFY10a | Os03g0402800 | 3.0 | 1 |
|  | OsJAZ8/9/TIFY10c | Os09g0439200 | 2.5 | 3 |
|  | OsJAZ2/TIFY11c-like | Os03g0180900 | 3.7 | - |
| α-Linolenic acid metabolism | OsLOX1a | Os03g0738600 | 3.0 | - |
|  | OsLOX1b | Os03g0700700 | 2.4 | 2 |
|  | OsLOX2 | Os08g0508800 | 2.4 | - |
|  | OsOPR1 | Os06g0216300 | 5.1 | 1 |
|  | OsOPR6 | Os06g0215500 | 7.6 | - |
| Auxin signaling | OsGH3-6 | Os05g0143800 | 2.3 | - |
|  | OsGH3-8 | Os07g0592600 | 2.1 | 5 |
| WRKY genes | OsWRKY8 | Os05g0583000 | 6.7 | 1 |
|  | OsWRKY108 | Os01g0821300 | 5.5 | 1 |
|  | OsWRKY24 | Os01g0714800 | 5.2 | 1 |
|  | OsWRKY76 | Os09g0417600 | 4.5 | 1 |
|  | OsWRKY67 | Os11g0117400 | 3.6 | - |
|  | OsWRKY113 | Os06g0158100 | 2.5 | - |
|  | OsWRKY62 | Os09g0417800 | 2.2 | - |
|  | OsWRKY15 | Os01g0656400 | 2.0 | 1 |
| Ethylene signaling (ERF genes) | OsERF67 | Os07g0674800 | 7.8 | 2 |
|  | OsERF118 | Os11g0168500 | 7.6 | 2 |
|  | OsERF27 | Os02g0676800 | 5.1 | - |
|  | OsERF109a | Os02g0764700 | 4.6 | 2 |
|  | OsERF108 | Os02g0781300 | 4.2 | - |
|  | OsERF109b | Os08g0474000 | 2.5 | 1 |
| Other transcription factors | ONAC59 | Os01g0862800 | 6.1 | 2 |
|  | bHLH148-like | Os01g0773800 | 8.9 | - |
| Dehydrin | OsLEA22 | Os01g0702500 | 9.0 | - |
|  | OsLEA2 | Os01g0314800 | 2.0 | 1 |
| Universal Stress Protein | OsUSP36 | Os10g0437500 | 5.2 | 1 |
| Chaperon proteins | Chaperon protein dnaJ | Os01g0702450 | 4.2 | - |
|  |  | Os08g0452900 | 3.6 | 1 |
| Early Response Dehyd. | OsERD7, chloroplastic | Os03g0241900 | 3.0 | 3 |
| Trehalose biosynthesis | Trehalose-6-phosphate synthase | Os08g0445700 | 3.6 | 1 |
|  |  | Os08g0414700 | 2.3 | - |
| Lignin biosynthesis | OsCCR | Os08g0277200 | 6.2 | 1 |
|  | OsC4H | Os02g0467000 | 6.1 | 1 |
| Terpene synthesis | OsGGPS7 | Os01g0248701 | 4.8 | - |
|  | OsTPS3 | Os02g0121700 | 5.4 | - |
|  | OsTPS23 | Os04g0344400 | 6.3 | - |
| Polyamine catabolism | OsPOA4 | Os04g0671200 | 3.7 | 3 |
|  | OsPOA6 | Os09g0368200 | 4.6 | - |

### Auxin signaling

Besides JA signaling, other plant hormone signaling pathways are also likely involved in DREB1a mechanism. Most notably, *GH3-6* and *GH3-8* that encode indole-3-acetic acid (IAA)-amido synthetases are upregulated in T_CS (**Table 2**). These enzymes catalyze conjugation of IAA with amino acids leading to its depletion (Staswick et al., 2005). Thus, upregulation of *GH3-6* and *GH3-8* in T_CS likely leads to attenuation of IAA signaling. Previous studies showed *GH3* induction during stress conditions including cold and drought (Han et al., 2020; Du et al., 2012). While many *GH3* are induced by ABA, it is not clear whether *GH3-6* and *GH3-8* are ABA-dependent. Further, while the promoter of *OsGH3-6* lacks *DRE* elements, that of *OsGH3-8* (Os07g0592600) contains 5 *DRE* elements (**Table 2**). Thus, *OsGH3-8* could be a direct target of DREB1a. The function of *OsGH3-8* in abiotic stress tolerance has not been demonstrated so far but Ding et al. (2008) showed that *OsGH3-8* enhances resistance to rice pathogen. Further, Zhang et al. (2009) showed that induction of *GH3* not only attenuates IAA signaling but also induces expression of late embryogenesis abundant (*LEA*) genes. Many *LEA* genes are induced during desiccation stress including freezing stress. Accordingly, *OsLEA22* was ~9 fold upregulated in T_CS compared to N_CS (**Table 2**). *OsLEA22* gene promoter lacks *DRE* elements; therefore, its upregulation is likely mediated through modulation of IAA signaling.

### *WRKY* transcription factors

Rice contains a large family of *WRKY* genes, majority of which is uncharacterized so far. Initially, *WRKY* were described to have a role in insect and pathogen response but their role in abiotic stress is becoming increasingly evident (Rushton et al., 2010; Viana et al., 2018). Eight *WRKY* TFs were upregulated in T_CS, five of which contained 1 *DRE* element each (*OsWRKY-8, −15, −24, −76, −108*) (**Table 2**). Of these *OsWRKY76*, a transcriptional repressor, has been characterized as having a direct role in cold tolerance (Yokotani et al., 2013). Although, it is not clear how *OsWRKY76* improves cold tolerance, it could be involved in repressing the negative regulators of cold-response genes. In addition, *OsWRKY −8, −24*, and −*108* are also upregulated >4-fold in T_CS and are likely involved in DREB1a-mediated abiotic stress tolerance.

### Other transcription factors

Besides *WRKY*, other TFs were also upregulated in T_CS. These include ethylene responsive factors (*ERFs*), basic helix loop helix (*bHLH*), and *NAC* (NAM ATAF1/2 CUC2) (**Table 2**). Of these, *ERFs* and *NAC* are well-known to positively regulate abiotic stress tolerance including cold stress (Hoang et al., 2017). Thus, multiple TFs that regulate abiotic stress are significantly upregulated in T_CS, suggesting that DREB1a regulates a complex network of genes associated with cold tolerance.

### Protective molecules

*OsLEA22* was ~9-fold upregulated and *OsLEA9* was ~2-fold upregulated in T_CS. LEA are extremely hydrophilic proteins that protect cellular membranes and proteins from desiccation injury (Bray, 1993; Cuming, 1999). Notably, LEA proteins cooperate with trehalose in this process (Goyal et al., 2005). Accordingly, two trehalose 6 phosphate synthase (*OsTPS*) genes were 2-fold and 3.5-fold upregulated in T_CS (**Table 2**). Next, *OsUSP36* was >5-fold upregulated in T_CS. USP36 is a member of universal stress proteins (USP) that is induced by cold stress (Arabia et al., 2021). The precise mechanism by which USPs bring about stress tolerance varies between USPs, but *OsUSP36* could be a target of DREB1a as its promoter includes a *DRE* element. Further, two genes encoding chaperon proteins dnaJ were also markedly upregulated in T_CS (~4-fold), possibly providing protective function to cellular components (**Table 2**). Finally, rice *Early Response to Dehydration7 (ERD7)* was upregulated 3-fold in T_CS. Arabidopsis ERD7 accumulates in response to multiple abiotic stresses (Cheng et al., 2013; Rasmussen et al., 2013). It is unclear how *ERD7* overexpression improves abiotic stress tolerance. However, a recent study reported its association with phospholipids and membranes and the susceptibility of *erd7* mutant to cold stress (Barajas-Lopez et al., 2021), suggesting a role of ERD7 in protecting cellular membranes from stress induced damage.

### Secondary metabolites

Plants synthesize a wide range of secondary metabolites that accumulate in response to environmental stimuli. Abundance of literature shows that secondary metabolites serve a protective function during stress (reviewed by Edreva et al., 2008; Wang et al., 2018; Yang et al., 2018). Several genes involved in secondary metabolite production were markedly upregulated in T_CS that include genes encoding Geranylgeranyl pyrophosphate synthase (GGPS), terpene synthases (TPS), phenylpropanoid (lignin) pathway, and polyamine oxidase (PAO). *OsGGPS, OsTPS3* and *OsTPS23* were upregulated 4 – 5-fold in T_CS, suggesting a role for terpenoids in DREB1a mediated abiotic stress tolerance. Jung et al. (2021) found induction of *OsTPS3* as part of the mechanism associated with improved drought tolerance. Phenylpropanoid pathway genes, *OsCCR* and *OsC4H* that encode Cinnamoyl-CoA reductase and Cinnamate 4-hydroxylase, respectively, were upregulated > 6-fold in T_CS (**Table 2**). These genes are involved in lignin biosynthesis, and lignin is a well-known cell wall biopolymer that imparts wall strength (Vanholme et al., 2010). Thus, DREB1a overexpressing plants possibly adapt to the stress by making their cell wall more rigid and impervious through lignification. Rice genome contains 7 polyamine oxidase (*OsPAO*) genes that are involved in back-conversion of polyamines towards maintaining polyamine homeostasis during stress (Ono et al., 2012; Yu et al., 2019). *OsPAO4* and *OsPAO6* were upregulated 3 - 4 fold in T_CS (**Table 2**). Sagor et al. (2021) found that *OsPAO4* and *OsPAO6* are rapidly induced by abiotic stresses including cold, salinity, drought, and wounding. In fact, *OsPAO6* was found to be massively induced by salinity stress. The catalytic activity of POA leads to H_2_O_2_ production that presumably functions as a signal molecule in inducing defense pathways (Angilini et al., 2010; Gupta et al., 2016).

### Environment sensing and signal transduction

Protein phosphorylation, cell communication, signal transduction, and chemical response were among GO Biological Processes associated with upregulated genes in T_CS, and plasma membrane, cell periphery, and intrinsic component of plasma membrane were associated with GO Cellular Component (**Table 1**). Accordingly, a number of receptor like kinases (RLK) were markedly upregulated to 4 - 7 fold. For example, wall associated receptor like kinase (*OsDEES1*; Os09g056160), Ser/Thr receptor like kinase (*OsRLCK253a*; Os08g0374600), putative leucine-rich repeat (LRR) receptor-like protein kinase At2g19210 (Os05g0524500), serine/threonine-protein kinase NAK (Os04g0563900), LRR receptor-like serine/threonine-protein kinase GSO2 (Os01g0162200). Additionally, Ca++ sensing proteins such as calmodulin (*OsCAM1*; Os03g0319300) and calcium dependent protein kinase (*CDPF*; Os08g0540400) were upregulated to 2.0 and 4.1-fold, respectively. Over-representation of these genes in T_CS suggests early sensing of the environmental stress and efficient signal transduction that possibly leads to improved stress tolerance in DREB1a expressing rice plants.

### Rice DREB genes

Rice DREB1 gene family (*OsDREB1*) is upregulated in the early phase (2 - 6 hours) of the cold stress at 4 - 10 ^o^C (DasGupta et al., 2020; Doubouzet et al., 2003; Pradhan et al., 2019; Yun et al., 2010). To check if *OsDREB* genes play a synergistic effect in stress tolerance displayed by *RD29a:DREB1a* transgenic plants, normalized read counts for *OsDREB1* and *OsDREB2* genes were compared between genotypes and treatment (**Figure S2**). Although *OsDREB1* gene family has been described to express similarly in response to cold stress (Mao & Chen, 2012); in our study, only *OsDREB1C* (Os06g0127100) and *OsDREB1G* (Os02g0677300) were upregulated either by cold treatment in non-transgenic line (N_CS) or in the transgenic line irrespective of the treatment (T_RT and T_CS), suggesting a positive effect of Arabidopsis DREB1a on these *OsDREB1* genes. Other *OsDREB1* genes such as *OsDREB1a, OsDREB1B*, and *OsDREB1H* were cold induced in both genotypes, and *OsDREB2* paralogs generally remained unaltered between the two genotypes or by the treatment (**Figure S2**). *OsDREB1C* and *OsDREB1G* are significantly upregulated by drought, cold, and flooding tolerance (Mittal et al., 2012; Mohanty, 2021; Wang et al., 2011), and found to be highly expressed in drought or cold tolerant rice lines (Chawade et al., 2013; Jung et al., 2017). A recent study demonstrated a direct role of *OsDREB1G* in cold tolerance by developing *OsDREB1G* over-expressor rice (Japonica cv. Dongjin) that showed tolerance to cold but not to drought or salinity (Moon et al., 2019). Search of *DRE* element in the promoter regions of *OsDREB1C* and *OsDREB1G* showed that *OsDREB1G* contains 3 copies of DRE motif ACCGAC. Arabidopsis DREB1a binds more efficiently to this motif than OsDREB1a, providing an explanation for higher induction of *OsDREB1G* in DREB1a transgenic rice as compared to the cold-treated non-transgenic rice. Overall, induction of *OsDREB1C* and *OsDREB1G* even at room temperature possibly through the leaky expression of Arabidopsis DREB1a in our transgenic line could contribute to stress tolerance trait.

### Down-regulated genes

Gene Ontology analysis and a survey of the top downregulated genes in T_CS in comparison to N_CS showed that genes associated with response to heat such as heat-shock proteins and heat shock factors were significantly down-regulated in T_CS. Further, photosynthesis related genes such as Protochlorophyllide reductase, Chlorophyll a/b-binding proteins, and photosystem II were down-regulated in T_CS (**Figure S1**). One of the down-regulated heat-shock factors is *OsHsfB2b* (Os08g0546800), a Class B heat shock factor, found to negatively regulate drought and salt stress in rice (Xiang et al., 2013). Down-regulation of heat response and photosynthesis genes was not found in T_RT and N_RT or N_CS and N_RT comparisons. Therefore, the set of down-regulated genes in T_CS possibly reflects DREB1a-mediated reprogramming of the cellular response by not only activating defense but also curbing processes that interfere in the stress response.

### RNA-seq data validation

To confirm RNA-seq data, quantitative PCR analysis was conducted on the upregulated DEGs which contained *DRE* elements with following criteria 1) DEGs co-expressed under abiotic stress conditions, 2) TF genes co-expressed under abiotic stress conditions, and 3) possible direct targets of DREB1a but not necessarily co-expressed. The first category included three genes *OsRBOH* (Os12g0541300), *OsJAZ5* (Os03g0402800), and *OsJAZ11* (Os04g0395800). The second category involved four transcription factors *OsDREB1a* (Os09g0522200), *OsDERF5* (Os027g0764700), *ONAC059* (Os01g0862800), *OsbHLH148*, and *OsWRKY76* (Os09g0417600), and the third category included *OsFLS2* (Os04g0618700), *MAPKKK17_18* (Os05g0545300), OsCAM1 (Os03g0319300). This analysis found a good correlation between RNA-seq and qPCR (R^2^=0.85) and generally validated RNA-seq data (**Figure S3**).

## DISCUSSION

Multi-gene stacking via Cre-*lox* recombination generated heritable expression of a stacked locus that contained Arabidopsis *DREB1a* regulated by *RD29a* promoter along with 4 marker genes (Pathak & Srivastava, 2020). Arabidopsis DREB1a is a well-known master regulator of abiotic stress tolerance (Shinozaki & Yamaguchi-Shinozaki, 2000). We used *RD29a* promoter to express *DREB1a* as *RD29a* is abundantly induced within 3 – 6 hours of imposing abiotic stresses such as cold, osmotic, drought, and salt. More importantly, this expression is sustained for a longer time (>24 h) than *DREB1a* expression under the native promoter (See, Arabidopsis eFP Browser; Winter et al., 2007). Further, since *RD29a* is a direct target of DREB1a, a positive feedback regulation would occur under stress condition, potentially activating stress gene network above a threshold. Arabidopsis *DREB1a* is rapidly induced upon cold exposure with a gradual decline after 6 hours. Rice DREB1a (*OsDREB1a*) follows a similar expression pattern showing induction by cold as early as 40 min after exposure and declining after 24 h (Dubouzet et al., 2003). Thus, sustained expression of *DREB1a* by the feedback mechanism of *RD29a:DREB1a* gene could lead to extended stress-response, that in turn, could enhance stress tolerance.

Rice is a salinity sensitive crop in which the seedling is the most affected stage in the presence of salts (Shannon et al., 1998). One of the first observable response in plants after exposure to salinity stress is the reduction in shoot growth, which is a very sensitive indicator of stress (Claeys et al., 2014; Negrão et al., 2017). *RD29a:DREB1a* transgenic seedlings in our study showed greater shoot development under salinity stress compared to non-transgenic controls (**Figure 1a-c**). Under salinity stress, reduction in plant growth rate can be observed due to accumulation of Na^+^ ions and the inability of plants to absorb enough water that results in the decrease in the osmotic component of soil water potential (Munns, 1993; Munns &Tester, 2008; Tester & Davenport, 2003). Similarly, Muthurajan et al. (2021) observed 41% higher growth rate in in *RD29a:DREB1a* transgenic rice (*Oryza sativa* L. sub sp. indica) seedlings under NaCl stress than the non-transgenic lines. Accumulation of Na^+^ in leaf tissues results in necrosis of older leaves, starting at the tips and margins and working back through the leaf (Tester & Davenport, 2003). In our study, non-transgenic plants had a higher proportion of chlorotic and necrotic leaves under stress than transgenic plants (**Figure 1e-h**). Muthurajan et al. (2021) also found that *RD29a:DREB1a* transgenic rice under salinity stress showed delayed yellowing symptoms in leaves and reduced rate of chlorophyll degradation as compared to non-transgenic lines.

Rice plants are characterized as a semi-aquatic plant as they survive submergence for a certain period (Das & Uchimiya, 2002; Fukai & Cooper, 1995). Therefore, rice is sensitive to drought periods starting from seedling to reproductive stages. During the periods of drought, cellular dehydration affects several basic physiological processes such as reduction of leaf expansion, photosynthetic inhibition, leaf abscission, cell death and production of reactive oxygen species (ROS) (Taiz et al., 2017). One of the adaptative processes in plants subjected to water shortage is the acceleration of leaf senescence, reducing the cumulative water demand over the plant cycle and allowing recycle of scarce resources to reproductive stage (Pic et al., 2002). Normally the leaf death starts from leaf tips and extends to all parts of the plant and finally to all the tillers through processes involving chlorophyll degradation (Bunnag & Pongthai, 2013; Pic et al. 2002). In our study, *RD29a:DREB1a* transgenic plants showed symptoms of yellowing in leaves much later during water shortage than the non-transgenic plants (**Figure 2a**). Furthermore, after water recovery time the non-transgenic plants showed a higher percentage of chlorotic and necrotic leaves (**Figure 2b-d**), indicating that DREB1a confers tolerance to more severe drought in rice. These observations are consistent with previous studies that showed drought tolerance in vegetative stages for *indica* (Datta et al. 2012; Latha et al. 2019) and *japonica* (Oh et al. 2005) rice cultivars.

Stress perception leads to a complex network of gene expression. Important abiotic stresses such as drought, cold, salinity induce common gene networks including transcription factors, protein kinases, receptor-like kinases, hormone signaling, and genes associated with the production of osmoprotectants (DasGupta et al., 2020; Todaka et al., 2015). However, it is important to improve tolerance to drought, salinity, and cold in most cultivated rice varieties (Menguer et al., 2017). In other words, cold/drought/salinity induced gene regulatory network in the cultivated rice is arguably too weak (expression dosage) and/or too late (off-timing) to confer tolerance to sustained stress conditions. In this regard, *RD29a:DREB1a* gene emerges as an important tool for improving stress tolerance for long term stress condition. Stress signaling is often accompanied by phenotypic penalty. Constitutive overexpression of *DREB1a* has been reported to cause growth retardation in Arabidopsis, tobacco, tomato, and rice (Hsieh et al., 2002; Ito et al., 2006; Kasuga et al., 2004; Maruyama et al., 2004). However, conditional expression of *DREB1a* by *RD29a* promoter mitigates phenotypic effects (Kasuga et al., 1999; Ma et al., 2010) therefore, it is important to use inducible expression of *DREB1a* instead of constitutive expression.

Oh et al. (2005) expressed *DREB1a* using the maize ubiquitin-1 promoter in rice (cv. Nakdong) and observed improved tolerance to cold, salinity, and drought. Interestingly, they did not observe any growth retardation in rice, despite the strong constitutive expression of *DREB1a.* Using 14-days old seedlings for microarray analysis, Oh et al. found induction of a set of genes including *Dip1* or *Dehydrin1* (Os02g0669100), *Lip5* or *WSI724* (Os03g0655400), *Bowman Birk trypsin inhibitor2* (Os01g0127600), *Receptor kinase LRR repeats* (Os05g0486100), and *Protein Phosphatase 2C* (Os11g0242200). These genes are also upregulated >2-fold in our transgenic rice upon cold treatment (T_CS). In fact, induction levels of *Bowman Birk trypsin inhibitor2* and *Protein Phosphatase 2C* are 4.5 and 6.8, respectively, as compared to the cold-stressed non-transgenic controls (N_CS). Similarly, in *DREB1a*-expressing chrysanthemum, upregulation of dehydrin gene and a higher expression of *DREB1a* regulon in *RD29a:DREB1a* transgenic plants as compared to *35S:DREB1a* plants was inferred as the basis of higher tolerance to cold and drought stress (Ma et al., 2010). Ma et al. found higher upregulation of alcohol dehydrogenase (*ADH*) in *RD29a:DREB1a* plants. Consistent with their findings, we also found 2.2- and 3.3-fold upregulation of two *ADH* genes, Os11g0210500 and Os11g0210300, respectively, in T_CS.

Studying *DREB1a* regulon in 10-days old rice seedlings (cv. Nipponbare) expressing Arabidopsis or rice *DREB1a* under *35S* promoter, Ito et al. (2006) found upregulation of *α-amylase, Lip5, β-1,3-glucanase,* and *protease inhibitor* genes among others. These genes are also upregulated in our transgenic plants under cold stress. Notably, *β-1,3-glucanase* (Os01g0947000) was upregulated 7.5-fold and *α-amylase* (Os01g0357400) to 2.4-fold in T_CS as compared to the N_CS. Expression of these genes could lead to higher concentrations of soluble sugars that in turn could improve osmotic strength of the cell.

More recently, Latha et al. (2019) performed microarray on flag leaf of transgenic (*RD29a:DREB1a*) *indica* rice subjected to drought at the booting stage for 14 days. They found upregulation of a large set of genes associated with chloroplast structure and function and suggested enhanced photosynthesis during stress as part of the mechanism for transgenic rice to withstand stress. In our dataset, however, chloroplast organization and function were downregulated in T_CS. Gene set enrichment analysis using GAGE method (Luo et al., 2009) indicated downregulation of photosynthesis related genes in T_CS as compared to N_CS (**Figure S4**). This difference could be based on the stage of plants used for gene expression analysis. Latha et al. used flag leaves from plants entering reproductive stage, while we used 10 days old seedlings. Clearly their plants were much more photosynthesis active in the greenhouse compared to our seedlings grown on artificial media in relatively lower light intensity.

Although each study, including ours, looked at a snap-shot of transcriptome (single timepoint) in a single tissue type, similar genes such as dehydrins, protein kinases, osmoprotectants were found to be associated with DREB1a-mediated stress response. This suggests that DREB1a activates similar downstream targets at possibly different time points in different plant species and cultivars. Overall, our transgenic plants under cold stress also showed upregulation of well-known cold, drought or salt stress genes as described previously by others. However, a few differences are noteworthy. First, we observed enrichment of genes associated with JA signaling in T_CS in comparison to N_CS. JA signaling has recently been recognized as an important part of abiotic stress regulation (Kazan, 2015; Santino et al., 2013); however, it has not been reported in *DREB1a* overexpressing plants, so far. JAZ proteins are central to JA signaling, wherein JA marks JAZ repressors for degradation, unleashing TFs such as bHLH, MYC2 that regulate the expression of stress response genes including DREB1a (Kazan & Manners, 2013). Thus, in the genetic pathway of abiotic stress response, JAZs are upstream of DREB1a (Kazan, 2015). It is therefore surprising that JAZ regulon was significantly upregulated in T_CS. Whether JAZ upregulation in T_CS has any direct role in stress tolerance cannot be confirmed. However, it is possible that JAZ upregulation is a mechanism to counter JAZ depletion due to enhanced production of endogenous JA upon stress. This possibly keeps other stress pathways such as ABA-dependent pathway suppressed through JAZ –TF interaction and prevents a costly run-off stress response, an important part of sustainable stress tolerance and survival.

Second, we observed enrichment of *WRKY* TF genes in T_CS in comparison to N_CS. Numerous studies on different plant species, including rice, show that WRKY are both positive and negative regulator of stresses. Through a web of interactions, much of which is yet to be understood, WRKY regulate stress response in both ABA-dependent and independent manner. (Bakshi & Oelmüller, 2014; Hong et al., 2017; Phukan et al., 2016; Viana et al., 2018). Although first classified as the regulators of biotic stresses, it is becoming increasingly clear that WRKY play important roles in drought, salinity, and cold stress response through activation or repression of their target genes. It is therefore surprising that only one study, so far, described differential expression of *WRKY* TFs in *RD29a:DREB1a* transgenic plants (*Salvia miltiorrhiza*) (Wei et al., 2016). This is likely due to the transient expression of *WRKY* in a complex web of cross-regulation with other TFs and MAP kinases that reprograms transcriptional machinery to facilitate response to disparate stresses such as drought and biotic stress by a single *WRKY* (Rushton et al., 2010). It is also likely that majority of *WRKY* are involved in drought and salinity stress as more and more *WRKY* are found to be associated with drought and salinity stress response, and only a few such as *OsWRKY76* are associated with cold stress response (Yokotani et al., 2013). *OsWRKY76* was among the highly upregulated *WRKYs* in our study (**Table 2**). However, to confirm the function of WRKYs found in our study, additional investigations based on transgenic overexpression and targeted mutagenesis are needed.

Next, we observed a set of *OsERF* upregulated in T_CS in comparison to N_CS controls, four of which are upregulated >4-fold (**Table 2**). Of these, *OsERF67, OsERF27, OsERF109a*, and *OsERF109b* were reported to be highly upregulated by cold-stress in 7-days old seedlings of *indica* rice variety Pusa Basmati and in a cold-tolerant *japonica* rice variety Jumli Marshi (Chawade et al., 2013; Mittal et al., 2012). In addition to cold-stress, *OsERFs* have also been found to be associated with submergence and drought stress (Wang et al., 2011; Wu et al., 2020).

Finally, we also observed unique sets of down-regulated genes, most notably, heat response and photosynthesis genes in T_CS. Heat stress response is distinct from dehydration stress such as drought/salt/cold stress, and the expression of heat-shock proteins under heat is a unique tolerance mechanism against heat stress. Therefore, down-regulation of heat shock protein and heat shock factor genes in T_CS suggests that heat tolerance pathway is possibly antagonistic to dehydration tolerance pathway and suppression of cellular response to heat positively regulates response to dehydration stress. Accordingly, Xiang et al. (2013) found that overexpression of OsHsfB2b in rice conferred heat tolerance but also rendered the plants more sensitive to drought and salt stress. Next, down-regulation of photosynthesis genes was also unique to T_CS. Multiple Chlorophyl a/b binding protein genes among other photosynthesis genes were significantly down-regulated in T_CS in comparison to N_CS (**Figure S4**). This could suggest that DREB1a reprograms the cell by diverting the energy from primary metabolism to secondary metabolism that synthesizes numerous secondary metabolites that generate a protective environment for the cell.

In conclusion, expression of *RD29a:DREB1a* from a stacked locus consisting of genes expressed by constitutive and inducible promoters was effective in enhancing salinity and drought tolerance in the *japonica* rice Taipei-309. Global transcriptomic analysis of the transgenic plants showed a complex interaction of transcription factors and hormone signaling, including JA and auxin signaling, towards reprograming the plant’s metabolic and growth pattern and enhancing the production of protective proteins and secondary metabolites for combating the stress (**Figure 6**). Rapid induction of the stress-response through early sensing of the stress, abundant expression of the stress regulon, and suppression of antagonistic pathways is likely the basis of improved stress tolerance in transgenic rice expressing Arabidopsis DREB1a.

## Acknowledgements

This project was supported in part by NSF-EPSCoR grant 1826836 to VS.

## Contributions

BP and YVB carried out salinity experiments, YVB and FB designed and performed drought experiments, BP developed RNA-seq outputs, bioinformatics analysis pipeline, and qPCR analysis, CM carried out DRE search, YVB and FB analyzed phenotypic data, VS analyzed overall data and wrote the paper.

## REFERENCES

Afgan E, Baker D, Batut B, Van Den Beek M, Bouvier D, Čech M, Chilton J, Clements D, Coraor N, Grüning BA, and Guerler A (2018) The Galaxy platform for accessible, reproducible and collaborative biomedical analyses: 2018 update. Nucleic Acids Res. 46(W1):W537–W544. doi: 10.1093/nar/gky379

Almeida DM, Almadanim MC, Lourenço T, Abreu IA, Saibo NJ, and Oliveira, MM (2016). Screening for abiotic stress tolerance in rice: salt, cold, and drought. In: Environmental Responses in Plants. Humana Press, New York, NY. pp. 155–182. doi:10.1007/978-1-4939-3356-3_14

Anbazhagan K, Bhatnagar-Mathur P, Vadez V, Dumbala SR, Kishor PK, and Sharma KK (2015) DREB1A overexpression in transgenic chickpea alters key traits influencing plant water budget across water regimes. Plant Cell Rep. 34:199–210. doi: 10.1007/s00299-014-1699-z.

Angelini R, Cona A, Federico R, Fincato P, Tavladoraki P, and Tisi A (2010) Plant amine oxidases “on the move”: an update. Plant Physiol. Biochem. 48:560–4. doi: 10.1016/j.plaphy.2010.02.001

Arabia S, Sami AA, Akhter S, Sarker RH, and Islam T (2021) Comprehensive *in silico* characterization of Universal Stress Proteins in rice (*Oryza sativa* L.) with insight into their stress-specific transcriptional modulation. Front. Plant Sci. 12:712607. doi: 10.3389/fpls.2021.712607.

Bakshi M and Oelmüller R (2014) WRKY transcription factors: Jack of many trades in plants. Plant Signal Behav. 9(2):e27700. doi:10.4161/psb.27700

Barajas-Lopez J, Tiwari A, Zarza X, Shaw MW, Pascual J, Punkkinen M, Bakowska JC, Munnik T, and Fujii H (2021) EARLY RESPONSE TO DEHYDRATION 7 remodels cell membrane lipid composition during cold stress in Arabidopsis. Plant and Cell Physiol. 62:80–91. https://doi.org/10.1093/pcp/pcaa139

Behnam B, Kikuchi A, Celebi-Toprak F, Kasuga M, Yamaguchi-Shinozaki K, and Watanabe KN (2007) Arabidopsis rd29A::DREB1A enhances freezing tolerance in transgenic potato. Plant Cell Rep. 26:1275–82. doi:10.1007/s00299-007-0360-5

Bhalani H, Thankappan R, Mishra GP, Sarkar T, Bosamia TC, and Dobaria JR (2019) Regulation of antioxidant mechanisms by AtDREB1A improves soil-moisture deficit stress tolerance in transgenic peanut (Arachis hypogaea L.). PloS One. 14(5):e0216706. doi: 10.1371/journal.pone.0216706

Bhatnagar-Mathur P, Devi MJ, Reddy DS, Lavanya M, Vadez V, Serraj R, Yamaguchi-Shinozaki K, and Sharma KK (2007) Stress-inducible expression of AtDREB1A in transgenic peanut (Arachis hypogaea L.) increases transpiration efficiency under water-limiting conditions. Plant Cell Rep. 26:2071–82. doi: 10.1007/s00299-007-0406-8

Bray EA (1993) Molecular responses to water deficit Plant Physiol. 103: 1035–1040. https://doi.org/10.1104/pp.103.4.1035

Bunnag S and Pongthai P (2013). Selection of rice *(Oryza sativa* L.) cultivars tolerant to drought stress at the vegetative stage under field conditions. Am. J. Plant Sci. 4:1701. http://dx.doi.org/10.4236/ajps.2013.49207

Claeys H, Van Landeghem S, Dubois M, Maleux K, and Inzé D (2014) What is stress? Doseresponse effects in commonly used in vitro stress assays. Plant Physiol. 165:519–527. doi: 10.1104/pp.113.234641.

Chawade A, Lindlöf A, Olsson B, and Olsson O (2013) Global expression profiling of low temperature induced genes in the chilling tolerant japonica rice Jumli Marshi. PLoS One. 8(12):e81729. Doi:10.1371/journal.pone.0081729

Cheng MC, Liao PM, Kuo WW, and Lin TP (2013) The Arabidopsis ETHYLENE RESPONSE FACTOR1 regulates abiotic stress-responsive gene expression by binding to different cis-acting elements in response to different stress signals. Plant Physiol. 162:1566–82. doi: 10.1104/pp.113.221911.

Chini A, Fonseca S, Fernández G, Adie B, Chico JM, Lorenzo O, García-Casado G, López-Vidriero I, Lozano FM, Ponce MR, Micol JL, and Solano R (2007) The JAZ family of repressors is the missing link in jasmonate signalling. Nature. 448:666–71. doi: 10.1038/nature06006.

Chow CN, Lee TY, Hung YC, Li GZ, Tseng KC, Liu YH, Kuo PL, Zheng HQ, and Chang WC (2019) PlantPAN3.0: a new and updated resource for reconstructing transcriptional regulatory networks from ChIP-seq experiments in plants. Nucleic Acids Res. 47(D1):D1155–D1163. doi: 10.1093/nar/gky1081.

Conesa A, Götz S, García-Gómez JM, Terol J, Talón M, and Robles M (2005) Blast2GO: a universal tool for annotation, visualization and analysis in functional genomics research. Bioinformatics. 21:3674–6. doi: 10.1093/bioinformatics/bti610

Conesa A and Götz S (2008) Blast2GO: A comprehensive suite for functional analysis in plant genomics. Int. J. Plant Genom. 2008; 2008:619832. doi: 10.1155/2008/619832.

Cuming AC (1999) LEA Proteins. In: Shewry P.R., Casey R. (eds) Seed Proteins. Springer, Dordrecht. https://doi.org/10.1007/978-94-011-4431-5_32

Dasgupta P, Das A, Datta S, Banerjee I, Tripathy S, and Chaudhuri S (2020) Understanding the early cold response mechanism in IR64 indica rice variety through comparative transcriptome analysis. BMC Genomics. 21:425. doi: 10.1186/s12864-020-06841-2

Das A and Uchimiya H (2002) Oxygen stress and adaptation of a semi-aquatic plant: rice *(Oryza sativa)*. J. Plant Res. 115:315–320. https://doi.org/10.1007/s10265-002-0043-9

Datta K, Baisakh N, Ganguly M, Krishnan S, Yamaguchi Shinozaki K, and Datta SK (2012). Overexpression of Arabidopsis and rice stress genes’ inducible transcription factor confers drought and salinity tolerance to rice. Plant Biotech. J. 10:579–586. https://doi.org/10.1111Zj.1467-7652.2012.00688.x

de Paiva Rolla AA, de Fátima Corrêa Carvalho J, Fuganti-Pagliarini R, Engels C, do Rio A, Marin SR, de Oliveira MC, Beneventi MA, Marcelino-Guimarães FC, Farias JR, Neumaier N, Nakashima K, Yamaguchi-Shinozaki K, and Nepomuceno AL (2014) Phenotyping soybean plants transformed with rd29A:AtDREB1A for drought tolerance in the greenhouse and field. Transgenic Res. 23:75–87. doi: 10.1007Zs11248-013-9723-6.

Devi MJ, Bhatnagar-Mathur P, Sharma KK, Serraj R, Anwar SY, and Vadez V (2011) Relationships between transpiration efficiency and its surrogate traits in the rd29A: DREB1A transgenic lines of groundnut. J. Agron. Crop Sci. 197:272–83. Doi:10.1111Zj.1439-037X.2011.00464.x

Ding X, Cao Y, Huang L, Zhao J, Xu C, Li X, and Wang S (2008) Activation of the indole-3-acetic acid-amido synthetase GH3-8 suppresses expansin expression and promotes salicylate- and jasmonate-independent basal immunity in rice. Plant Cell. 20:228–40. doi: 10.1105/tpc.107.055657.

Du H, Wu N, Fu J, Wang S, Li X, Xiao J, and Xiong L (2012) A GH3 family member, OsGH3-2, modulates auxin and abscisic acid levels and differentially affects drought and cold tolerance in rice. J. Exp. Bot. 63(18):6467–80. doi: 10.1093/jxb/ers300.

Dubouzet JG, Sakuma Y, Ito Y, Kasuga M, Dubouzet EG, Miura S, Seki M, Shinozaki K, and Yamaguchi-Shinozaki K (2003), *OsDREB* genes in rice, *Oryza sativa* L., encode transcription activators that function in drought-, high-salt- and cold-responsive gene expression. Plant J. 33:751–763. https://doi.org/10.1046/j.1365-313X.2003.01661.x

Edgar R, Domrachev M, and Lash AE (2002) Gene Expression Omnibus: NCBI gene expression and hybridization array data repository. Nucleic Acids Res. 30:207–10. doi: 10.1093ZnarZ30.1.207.

Edreva, A, Velikova V, Tsonev T, Dagnon S, Gürel A, Aktaş L, and Gesheva E (2008) Stress-protective role of secondary metabolites: diversity of functions and mechanisms. Gen. Appl. Plant Physiol 34.1-2: 67–78.

Fukai S and Cooper M (1995) Development of drought-resistant cultivars using physiomorphological traits in rice. Field Crops Res. 40:67–86. https://doi.org/10.1016/0378-4290(94)00096-U

Ganguly M, Roychoudhury A, Sengupta DN, Datta SK, and Datta K (2020) Independent overexpression of OsRab16A and AtDREB1A exhibit enhanced drought tolerance in transgenic aromatic rice variety Pusa Sugandhi 2. J. Plant Biochem. Biotech. 29:503–517. https://doi.org/10.1007/s13562-020-00565-w

Ge SX, Son EW, and Yao R (2019) iDEP: an integrated web application for differential expression and pathway analysis of RNA-Seq data. BMC Bioinform. 19: 534 (2018). https://doi.org/10.1186/s12859-018-2486-6

Geda CK, Repalli SK, Dash GK, Swain P, and Rao GJN (2019) Enhancement of drought tolerance in rice through introgression of Arabidopsis *DREB1A* through transgenic approach. J Rice Res. 7:208. DOI: 10.4172Z2375-4338.1000208

Goyal K, Walton LJ, and Tunnacliffe A (2005). LEA proteins prevent protein aggregation due to water stress. Biochem. J. 388(Pt 1):151–157. https://doi.org/10.1042/BJ20041931

Gupta K, Sengupta A, Chakraborty M, and Gupta B (2016) Hydrogen peroxide and polyamines act as double edged swords in plant abiotic stress responses. Front. Plant Sci. 7:1343. Doi:10.3389/fpls.2016.01343

Han B, Ma X, Cui D, Wang Y, Geng L, Cao G, Zhang H, and Han L (2020) Comprehensive evaluation and analysis of the mechanism of cold tolerance based on the transcriptome of weedy rice seedlings. Rice (N Y). 13(1):12. Doi:10.1186/s12284-019-0363-1

Hoang XLT, Nhi DNH, Thu NBA, Thao NP, and Tran LP (2017) Transcription factors and their roles in signal transduction in plants under abiotic stresses. Curr. Genom. 18:483–497. Doi:10.2174/1389202918666170227150057

Hong C, Cheng D, Zhang G, Zhu D, Chen Y, and Tan M (2017) The role of ZmWRKY4 in regulating maize antioxidant defense under cadmium stress. Biochem. Biophys. Res. Commun. 482:1504–1510. doi: 10.1016/j.bbrc.2016.12.064

Hong B, Tong Z, Ma N, Li J, Kasuga M, Yamaguchi-Shinozaki K, and Gao J (2006) Heterologous expression of the AtDREB1A gene in chrysanthemum increases drought and salt stress tolerance. Sci. China C Life Sci. 49:436–45. doi: 10.1007/s11427-006-2014-1.

Hsieh TH, Lee JT, Charng YY, and Chan, MT (2002) Tomato plants ectopically expressing Arabidopsis CBF1 show enhanced resistance to water deficit stress. Plant Physiol. 130: 618–626. https://doi.org/10.1104/pp.006783

Ito Y, Katsura K, Maruyama K, Taji T, Kobayashi M, Seki M, Shinozaki K, and Yamaguchi-Shinozaki K (2006) Functional analysis of rice DREB1/CBF-type transcription factors involved in cold-responsive gene expression in transgenic rice. Plant Cell Physiol. 47:141–53. doi: 10.1093/pcp/pci230.

Jung H, Chung PJ, Park SH, Redillas MCFR, Kim YS, Suh JW, and Kim JK (2017) Overexpression of OsERF48 causes regulation of OsCML16, a calmodulin-like protein gene that enhances root growth and drought tolerance. Plant Biotech. J. 15:1295–1308. doi: 10.1111/pbi.12716.

Jung SE, Bang SW, Kim SH, Seo JS, Yoon HB, Kim YS, and Kim JK (2021) Overexpression of OsERF83, a vascular tissue-specific transcription factor gene, confers drought tolerance in rice. Int. J. Mol. Sci. 22:7656. Doi:10.3390/ijms22147656

Kanehisa M, Sato Y, and Morishima K (2016) BlastKOALA and GhostKOALA: KEGG tools for functional characterization of genome and metagenome sequences. J. Mol. Biol. 428:726–731. doi: 10.1016/j.jmb.2015.11.006.

Kasuga M, Liu Q, Miura S, Yamaguchi-Shinozaki K, and Shinozaki K (1999) Improving plant drought, salt and freezing tolerance by gene transfer of a single stress-inducible transcription factor. Nat Biotechnol. 17:287–291. https://doi.org/10.1038/7036

Kasuga M, Miura S, Shinozaki K, and Yamaguchi-Shinozaki K (2004) A combination of the Arabidopsis DREB1A gene and stress-inducible rd29A promoter improved drought- and low-temperature stress tolerance in tobacco by gene transfer. Plant Cell Physiol. 45:346–50. doi: 10.1093/pcp/pch037.

Kazan K (2015) Diverse roles of jasmonates and ethylene in abiotic stress tolerance. Trends Plant Sci. 20: 219–229. https://doi.org/10.1016/j.tplants.2015.02.001.

Kazan K and Manners JM (2013) MYC2: the master in action. Mol. Plant. 6:686–703. https://doi.org/10.1093/mp/sss128.

Kudo M, Kidokoro S, Yoshida T, Mizoi J, Kojima M, Takebayashi Y, Sakakibara H, Fernie AR, Shinozaki K, and Yamaguchi-Shinozaki K (2019) A gene-stacking approach to overcome the trade-off between drought stress tolerance and growth in Arabidopsis. Plant J. 97:240–256. doi: 10.1111/tpj.14110.

Latha GM, Raman KV, Lima JM, Pattanayak D, Singh AK, Chinnusamy, V, Bansal KC, Rao KRSS, and Mohapatra T (2019) Genetic engineering of indica rice with AtDREB1A gene for enhanced abiotic stress tolerance. Plant Cell Tiss. Organ Cult. (PCTOC), 136:173–188. https://doi.org/10.1007/s11240-018-1505-7

Li X, Cao K, Wang C, Sun ZW, and Yan, L (2010) Variation of photosynthetic tolerance of rice cultivars (Oryza sativa L.) to chilling temperature in the light. African J. Biotech. 9:1325–1337. https://doi.org/10.5897/AJB10.922

Li H and Durbin R (2010) Fast and accurate long-read alignment with Burrows-Wheeler transform. Bioinformatics. 26:589–95. doi: 10.1093/bioinformatics/btp698.

Liao Y, Smyth GK, and Shi W (2014) featureCounts: an efficient general purpose program for assigning sequence reads to genomic features. Bioinformatics. 30:923–30.

Livak KJ and Schmittgen TD (2001) Analysis of relative gene expression data using real-time quantitative PCR and the 2(-Delta Delta C(T)) method. Methods. 25(4):402–8. doi: 10.1006/meth.2001.1262.

Liu R, Holik AZ, Su S, Jansz N, Chen K, Leong HS, Blewitt ME, Asselin-Labat ML, Smyth GK, and Ritchie ME (2015) Why weight? Modelling sample and observational level variability improves power in RNA-seq analyses. Nucleic Acids Res. 43(15):e97. doi: 10.1093/nar/gkv412.

Luo W, Friedman MS, Shedden, K, Hankenson KD, and Woolf PJ (2009) GAGE: generally applicable gene set enrichment for pathway analysis. BMC Bioinformatics 10:161 https://doi.org/10.1186/1471-2105-10-161

Ma C, Hong B, Wang T, Yang JY, Tong Z, Zuo ZR, Yamaguchi-Shinozaki K, and Gao JP (2010) DREB1A regulon expression in *rd29A:DREB1A* transgenic chrysanthemum under low temperature or dehydration stress. J. Hort. Sci. Biotech., 85:6, 503–510, doi: 10.1080/14620316.2010.11512705

Mao D and Chen C (2012) Colinearity and similar expression pattern of rice *DREB1s* reveal their functional conservation in the cold-responsive pathway. PLoS ONE 7(10): e47275. https://doi.org/10.1371/journal.pone.0047275

Maruyama K, Sakuma Y, Kasuga M, Ito Y, Seki M, Goda H, Shimada Y, Yoshida S, Shinozaki K, and Yamaguchi-Shinozaki K (2004) Identification of cold-inducible downstream genes of the Arabidopsis DREB1A/CBF3 transcriptional factor using two microarray systems. Plant J. 38: 982–993. DOI: 10.1111/j.1365-313X.2004.02100.x

Menguer PK, Sperotto RA, and Ricachenevsky FK (2017). A walk on the wild side: Oryza species as source for rice abiotic stress tolerance. Genet. Mol. Biol. 40(1 suppl 1):238–252. Doi:10.1590/1678-4685-GMB-2016-0093

Mittal D, Madhyastha DA, and Grover A (2012) Genome-wide transcriptional profiles during temperature and oxidative stress reveal coordinated expression patterns and overlapping regulons in rice. PLoS ONE 7(7): e40899. https://doi.org/10.1371/journal.pone.0040899

Mohanty B (2021) Promoter architecture and transcriptional regulation of genes upregulated in germination and coleoptile elongation of diverse rice genotypes tolerant to submergence. Front. Genet. 12:639654. Doi:10.3389/fgene.2021.639654

Moldenhauer K, Counce P, and Hardke J (2018) Rice growth and development. Rice Production Handbook, 192, 7–14. https://www.uaex.uada.edu/publications/pdf/mp192/chapter-2-word.pdf

Moon SJ, Min MK, Kim JA, Kim DY, Yoon IS, Kwon TR, Byun MO, and Kim BG (2019) Ectopic expression of *OsDREB1G*, a member of the OsDREB1 subfamily, confers cold Stress tolerance in rice. Front. Plant Sci. 10:297. doi: 10.3389/fpls.2019.00297.

Msanne J, Lin J, Stone JM, and Awada T (2011) Characterization of abiotic stress-responsive Arabidopsis thaliana RD29A and RD29B genes and evaluation of transgenes. Planta. 234:97–107. doi: 10.1007/s00425-011-1387-y

Munns R (1993) Physiological processes limiting plant growth in saline soils: some dogmas and hypotheses. Plant Cell & Environ. 16:15–24. https://doi.org/10.1111/j.1365-3040.1993.tb00840.x

Munns R and Tester M (2008) Mechanisms of salinity tolerance. Annu. Rev. Plant Biol 59: 651–681. https://doi.org/10.1146/annurev.arplant.59.032607.092911

Muthurajan R, Ramanathan V, Bansilal Shillak A, Madhuri Pralhad S, Shankarrao CN, Rahman H, Kambale R, Nallathambi J, Tamilselvan S, and Madasamy P (2021) Controlled over-expression of AtDREB1A enhances tolerance against drought and salinity in rice. Agronomy 11(1):159. https://doi.org/10.3390/agronomy11010159

Negrão S, Schmöckel SM, and Tester M (2017). Evaluating physiological responses of plants to salinity stress. Annals Bot., 119:1–11. https://doi.org/10.1093/aob/mcw191

Oh SJ, Song SI, Kim YS, Jang HJ, Kim SY, Kim M, Kim YK, Nahm BH, and Kim JK (2005) Arabidopsis CBF3/DREB1A and ABF3 in transgenic rice increased tolerance to abiotic stress without stunting growth. Plant Physiol. 138:341–351. doi: 10.1104/pp.104.059147.

Ono Y, Kim DW, Watanabe K, Sasaki A, Niitsu M, Berberich T, Kusano T, and Takahashi Y (2012) Constitutively and highly expressed Oryza sativa polyamine oxidases localize in peroxisomes and catalyze polyamine back conversion. Amino Acids. 42:867–76. doi: 10.1007/s00726-011-1002-3.

Pathak B and Srivastava V (2020) Recombinase-mediated integration of a multigene cassette in rice leads to stable expression and inheritance of the stacked locus. Plant Direct. 4: 1–10. https://doi.org/10.1002/p1d3.236

Phukan UJ, Jeena GS, and Shukla RK (2016) WRKY Transcription Factors: Molecular regulation and stress responses in plants. Front. Plant Sci. 7:760; doi: 10.3389/fpls.2016.00760

Pic E, Serve, BT de la, Tardieu F, and Turc O (2002) Leaf senescence induced by mild water deficit follows the same sequence of macroscopic, biochemical, and molecular events as monocarpic senescence in pea. Plant Physiol. 128:236–246.https://doi.org/10.1104/pp.010634

Pradhan SK, Pandit E, Nayak DK, Behera L, and Mohapatra T (2019) Genes, pathways and transcription factors involved in seedling stage chilling stress tolerance in indica rice through RNA-Seq analysis. BMC Plant Biol. 19:352. doi: 10.1186/s12870-019-1922-8

R Core Team (2017) R: A language and environment for statistical computing. R Foundation for Statistical Computing. http://www.R-project.org. Accessed 20 Oct 2020.

Ravikumar G, Manimaran P, Voleti SR, Subrahmanyam D, Sundaram RM, Bansal KC, Viraktamath BC, and Balachandran SM (2014) Stress-inducible expression of AtDREB1A transcription factor greatly improves drought stress tolerance in transgenic indica rice. Transgenic Res. 23:421–39. doi: 10.1007Zs11248-013-9776-6.

Rasmussen S, Barah P, Suarez-Rodriguez MC, Bressendorff S, Friis P, Costantino P, Bones AM, Nielsen HB, and Mundy J (2013) Transcriptome responses to combinations of stresses in Arabidopsis. Plant Physiol. 161(4):1783–94. doi: 10.1104/pp.112.210773.

Reimand J, Arak T, Adler P, Kolberg L, Reisberg S, Peterson H, and Vilo J (2016) g:Profiler-a web server for functional interpretation of gene lists (2016 update). Nucleic Acids Res. 44(W1):W83–9. doi: 10.1093/nar/gkw199.

Rushton PJ, Somssich IE, Ringler P, and Shen QJ (2010) WRKY transcription factors. Trends Plant Sci. 15:247–258; https://doi.org/10.1016/j.tplants.2010.02.006.

SAS I. (2013) Base SAS 9.4 procedures guide: statistical procedures. Cary, NC, USA: SAS Institute Inc

Sagor GHM, Inoue M, Kusano T, and Berberich T (2021) Expression profile of seven polyamine oxidase genes in rice (*Oryza sativa*) in response to abiotic stresses, phytohormones and polyamines. Physiol. Mol. Biol. Plants. 27(6):1353–1359. doi: 10.1007/s12298-021-01006-1.

Santino A, Taurino M, De Domenico S, Bonsegna S, Poltronieri P, Pastor V, and Flors V (2013) Jasmonate signaling in plant development and defense response to multiple (a)biotic stresses. Plant Cell Rep. 32(7):1085–98. doi: 10.1007/s00299-013-1441-2.

Sarkar T, Thankappan R, Kumar A, Mishra GP, and Dobaria JR (2016) Stress inducible expression of AtDREB1A transcription factor in transgenic peanut (Arachis hypogaea L.) conferred tolerance to soil-moisture deficit stress. Front Plant Sci. 7:935. Doi:10.3389/fpls.2016.00935

Shailani A, Joshi R, Singla-Pareek SL, and Pareek A (2021) Stacking for future: Pyramiding genes to improve drought and salinity tolerance in rice. Physiol. Plant. 172:1352–1362. doi: 10.1111/ppl.13270.

Shannon M C, Rhoades JD, Draper JH, Scardaci SC, and Spyres MD (1998). Assessment of salt tolerance in rice cultivars in response to salinity problems in California. Crop Sci. 38:394–398. https://doi.org/10.2135/cropsci1998.0011183X003800020021x

Shimazaki T, Endo T, Kasuga M, Yamaguchi-Shinozaki K, Watanabe KN, and Kikuchi A (2016). Evaluation of the yield of abiotic-stress-tolerant AtDREB1A transgenic potato under saline conditions in advance of field trials. Breed Sci. 66:703–710. doi: 10.1270/jsbbs.16054.

Shinozaki K and Yamaguchi-Shinozaki K (2000). Molecular responses to dehydration and low temperature: differences and cross-talk between two stress signaling pathways. Curr. Opin. Plant Biol. 3:217–23. https://doi.org/10.1016/S1369-5266(00)80068-0

Smirnoff N and Bryant JA (1999) DREB takes the stress out of growing up. Nat Biotechnol. 17:229–30. Doi:10.1038/6968.

Srivastava V and Ow DW (2002) Biolistic mediated site-specific integration in rice. Mol. Breed. 8:345–349. https://doi.org/10.1023/A:1015229015022

Staswick PE, Serban B, Rowe M, Tiryaki I, Maldonado MT, Maldonado MC, and Suza W (2005) Characterization of an Arabidopsis enzyme family that conjugates amino acids to indole-3-acetic acid. Plant Cell 17:616–627. https://doi.org/10.1105/tpc.104.026690

Taiz L, Zeiger E, Moller IM, and Murphy A (2017) Fisiologia e desenvolvimento vegetal. 6.ed. Porto Alegre: Artmed, 2017. 858p.

Tester M and Davenport R (2003) Na+ tolerance and Na+ transport in higher plants. Ann. Bot. 91:503–527. https://doi.org/10.1093/aob/mcg058

Thines B, Katsir L, Melotto M, Niu Y, Mandaokar A, Liu G, Nomura K, He SY, Howe GA, and Browse J (2007) JAZ repressor proteins are targets of the SCF(COI1) complex during jasmonate signalling. Nature 448:661–5. doi: 10.1038/nature05960.

Todaka D, Shinozaki K, and Yamaguchi-Shinozaki K (2015) Recent advances in the dissection of drought-stress regulatory networks and strategies for development of drought-tolerant transgenic rice plants. Front. Plant Sci. 6:84. doi: 10.3389/fpls.2015.00084.

Vanholme R, Demedts B, Morreel K, Ralph J, and Boerjan W (2010) Lignin biosynthesis and structure. Plant Physiol. 153:895–905, https://doi.org/10.1104/pp.110.155119

Viana VE, Busanello C, da Maia LC, Pegoraro C, and Costa de Oliveira A (2018) Activation of rice WRKY transcription factors: an army of stress fighting soldiers? Curr. Opin. Plant Biol. 45(Pt B):268–275. Doi:10.1016/j.pbi.2018.07.00

Wang D, Pan Y, Zhao X, Zhu L, Fu B, and Li Z (2011) Genome-wide temporal-spatial gene expression profiling of drought responsiveness in rice. BMC Genomics. 12:149. Doi:10.1186/1471-2164-12-149

Wang W, Li Y, Dang P, Zhao S, Lai D, and Zhou, L. (2018). Rice Secondary Metabolites: Structures, roles, biosynthesis, and metabolic regulation. Molecules (Basel, Switzerland), 23:3098. https://doi.org/10.3390/molecules23123098

Wasternack C (2007) Jasmonates: an update on biosynthesis, signal transduction and action in plant stress response, growth and development. Ann. Bot. 100(4):681–97. doi: 10.1093/aob/mcm079.

Wei T, Deng K, Liu D, Gao Y, Liu Y, Yang M, Zhang L, Zheng X, Wang C, Song W, and Chen C (2016) Ectopic expression of DREB transcription factor, AtDREB1A, confers tolerance to drought in transgenic Salvia miltiorrhiza. Plant and Cell Physiol. 57:1593–609. doi: 10.1093/pcp/pcw084.

Winter D, Vinegar B, Nahal H, Ammar R, Wilson GV, and Provart NJ (2007) An “Electronic Fluorescent Pictograph” browser for exploring and analyzing large-scale biological data sets. PLoS ONE 2(8): e718. https://doi.org/10.1371/journal.pone.0000718

Wu YS and Yang CY (2020) Comprehensive transcriptomic analysis of auxin responses in submerged rice coleoptile growth. Int J. Mol Sci. 21:1292. Doi:10.3390/ijms21041292

Xiang J, Ran J, Zou J, Zhou X, Liu A, Zhang X, Peng Y, Tang N, Luo G, and Chen X (2013) Heat shock factor OsHsfB2b negatively regulates drought and salt tolerance in rice. Plant Cell Rep. 32:1795–806. doi: 10.1007/s00299-013-1492-4.

Yamaguchi-Shinozaki K and Shinozaki K (1994) A novel cis-acting element in an Arabidopsis gene is involved in responsiveness to drought, low-temperature or high-salt stress. Plant Cell 6:251–264. doi: 10.1105/tpc.6.2.251.

Yamaguchi-Shinozaki K and Shinozaki K (2006) Transcriptional regulatory networks in cellular responses and tolerance to dehydration and cold stresses. Annu. Rev. Plant Biol. 57:781–803. doi: 10.1146/annurev.arplant.57.032905.105444.

Yang L, Wen KS, Ruan X, Zhao YX, Wei F, and Wang Q (2018) Response of plant secondary metabolites to environmental factors. Molecules. 23:762. doi: 10.3390/molecules23040762.

Yokotani N, Sato Y, Tanabe S, Chujo T, Shimizu T, Okada K, Yamane H, Shimono M, Sugano S, Takatsuji H, Kaku H, Minami E, and Nishizawa Y (2013) WRKY76 is a rice transcriptional repressor playing opposite roles in blast disease resistance and cold stress tolerance. J. Exp. Bot. 64:5085–97. doi: 10.1093/jxb/ert298.

Yu Z, Jia D, and Liu T (2019) Polyamine Oxidases play various roles in plant development and abiotic stress tolerance. Plants (Basel) 8:184. Doi:10.3390/plants8060184

Yun KY, Park MR, Mohanty B, Herath V, Xu F, Mauleon R, Wijaya E, Bajic VE, Bruskiewich R, de los Reyes BG (2010) Transcriptional regulatory network triggered by oxidative signals configures the early response mechanisms of japonica rice to chilling stress. BMC Plant Biol 10, 16. https://doi.org/10.1186/1471-2229-10-16

Zhang H, Zhu J, Gong Z, and Zhu J-K (2021) Abiotic stress responses in plants. Nat Rev Genet https://doi.org/10.1038/s41576-021-00413-0

Zhang SW, Li CH, Cao J, Zhang Y-C, Zhang S-Q, Xia Y-F, Sun D-Y, and Sun Y (2009) Altered architecture and enhanced drought tolerance in rice via the down-regulation of indole-3-acetic acid by TLD1/OsGH3.13 activation. Plant Physiol. 151:1889–1901. Doi:10.1104/pp.109.146803

