## Supplementary material for "Phenotypic and transcriptomic analysis reveals early stress responses in transgenic rice expressing Arabidopsis DREB1a": Figure S1c

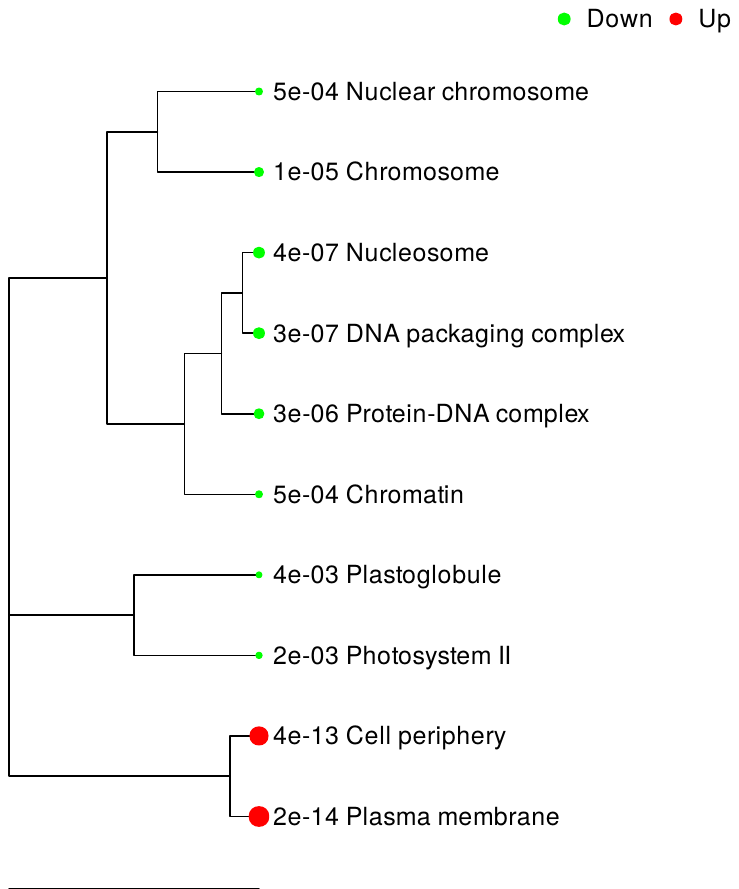

**GO: Cellular Component**

**Supplementary Fig. S1c:** Gene ontology (GO) analysis of enriched cellular components. Gene enrichment tree showing up- (red) or down-regulated (green) genes for GO: cellular components of 2069 DEGs (p-value ≤0.01) in cold-shocked *RD29a:DREB1a* transgenic lines in comparison to cold-shocked non-transgenic lines
