## Supplementary material for "Phenotypic and transcriptomic analysis reveals early stress responses in transgenic rice expressing Arabidopsis DREB1a": Figure S2

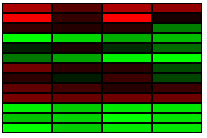

**OsDREB1A**

**OsDREB1B**

**OsDREB1C**

**OsDREB1D**

**OsDREB1E**

**OsDREB1F**

**OsDREB1G**

**OsDREB1H**

**OsDREB2A**

**OsDREB2A**

**OsDREB2A**

**OsDREB2A**

**OsDREB2E**

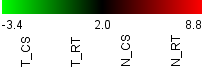

**Supplementary Fig. S2:** Relative expression of OsDREB1 and OsDREB2 regulon in RD29a:DREB1a transgenic (T) and non-transgenic (N) under cold-shock (CS) or room temperature (RT) control conditions. Normalized counts of mapped reads were used to generate the heatmap.
