## Supplementary material for "Phenotypic and transcriptomic analysis reveals early stress responses in transgenic rice expressing Arabidopsis DREB1a": Figure S3

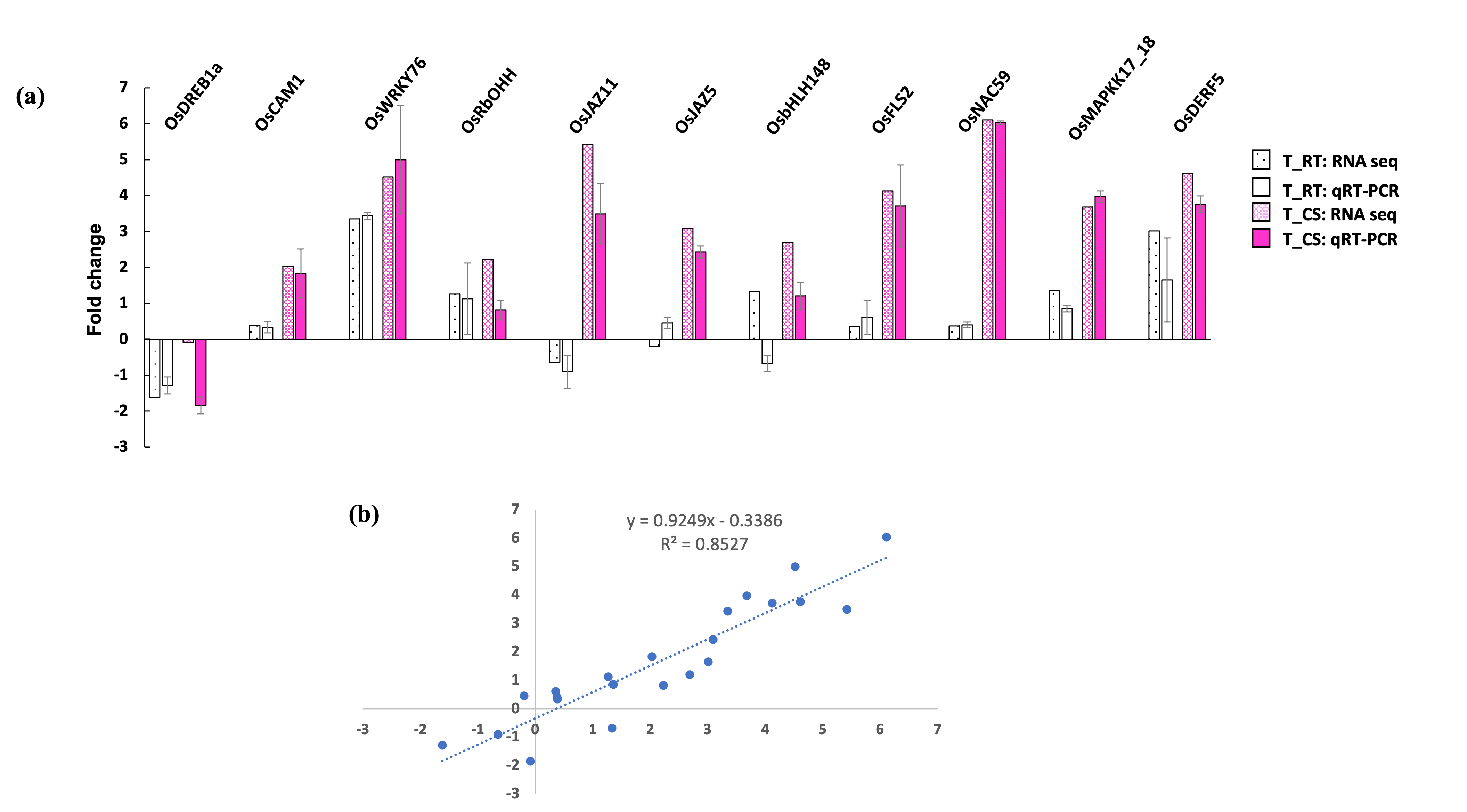

**Supplementary Fig. S3:** Validation of RNA-seq data by quantitative Reverse Transcriptase PCR (qRT-PCR). (**a**) Log2 fold change in the expression of selected genes in RD29a:DREB1a transgenic lines (T) at room temperature (RT) or upon cold-shock (CS) as determined by RNAseq or qRT-PCR. Fold change in gene expression was calculated relative to non-transgenic controls in respective treatments (RT or CS). (**b**) Correlation of RNAseq and qRT-PCR data (log2 fold change).
