## Supplementary material for "Phenotypic and transcriptomic analysis reveals early stress responses in transgenic rice expressing Arabidopsis DREB1a": Figure S4a

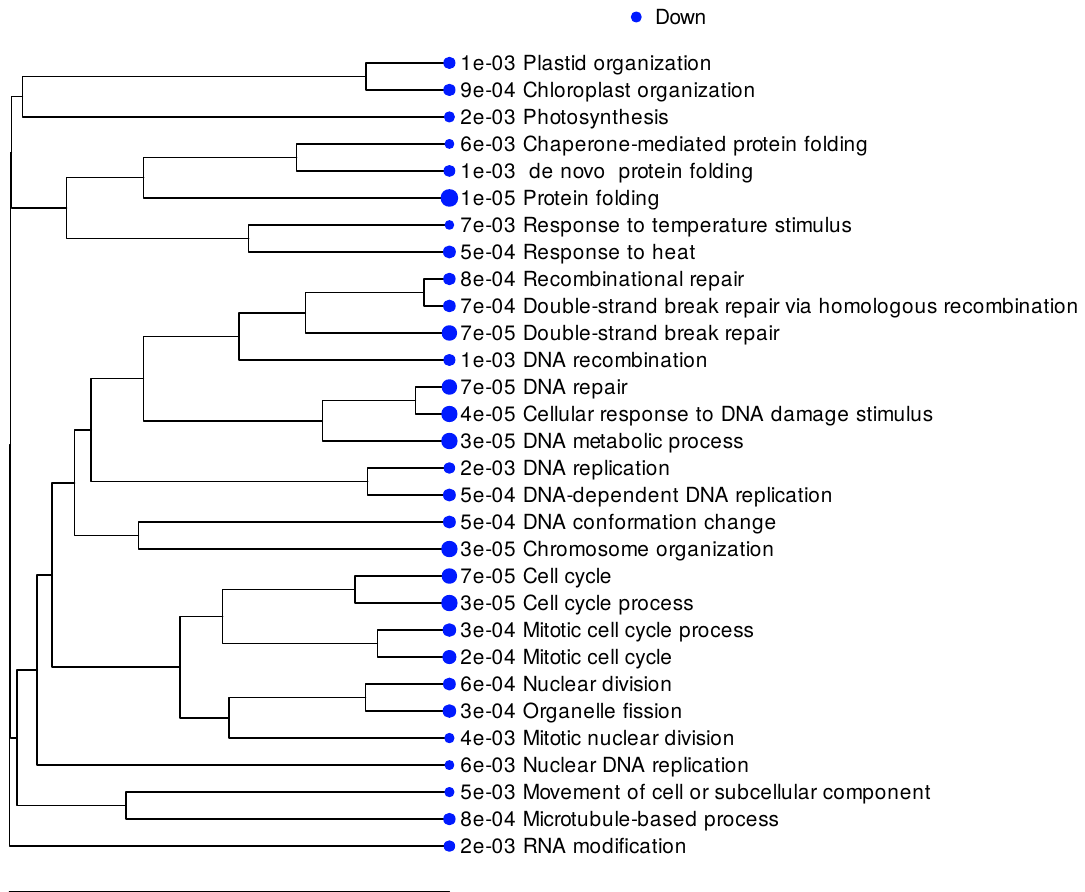

**Supplementary Fig. S4a:** Down-regulated biological processes in cold-shocked *RD29a:DREB1a* transgenic line in comparison to cold-shocked non-transgenic lines. Gene set enrichment analysis using GAGE method (Luo et al., 2009) for GO: biological processe. FDR= 0.01, Number of top pathways= 30. Data analyzed using iDEP.93v (http://bioinformatics.sdstate.edu/idep93/).
