## Supplementary material for "Phenotypic and transcriptomic analysis reveals early stress responses in transgenic rice expressing Arabidopsis DREB1a": Figure S4b

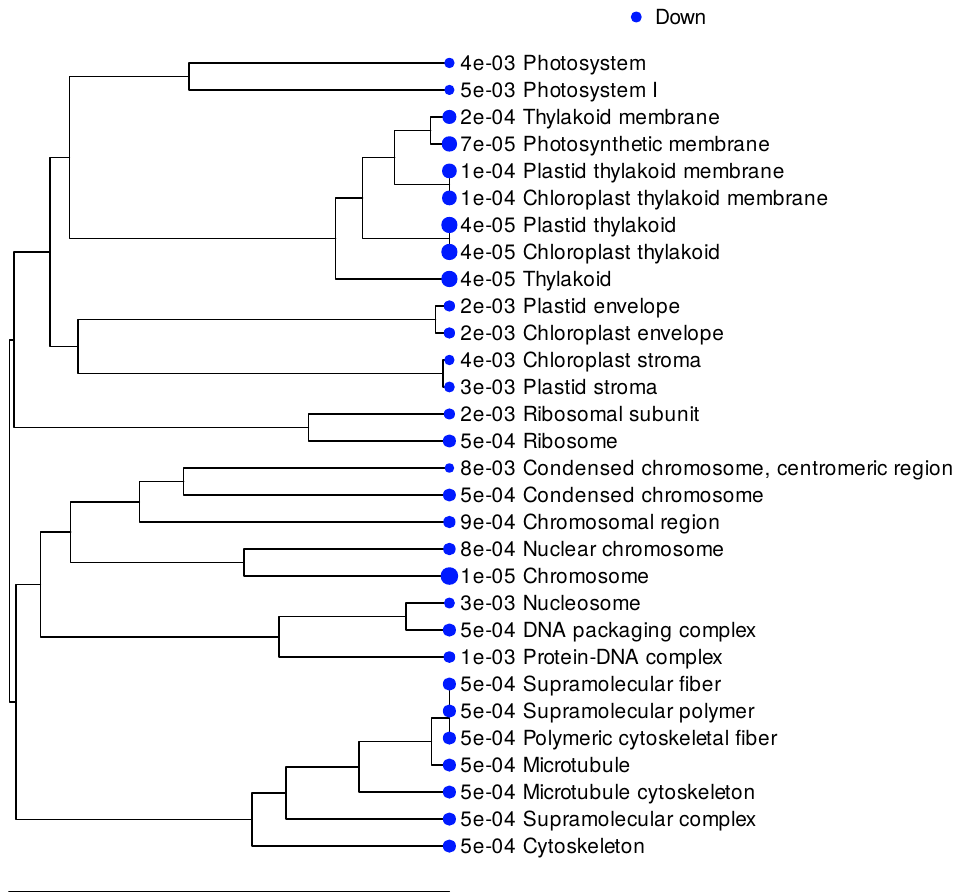

**Supplementary Fig. S4b:** Down-regulated molecular function in cold-shocked *RD29a:DREB1a* transgenic line in comparison to cold-shocked non-transgenic lines. Gene set enrichment analysis using GAGE method (Luo et al., 2009) for GO: molecular function. FDR= 0.01, Number of top pathways= 30. Data analyzed using iDEP.93v (http://bioinformatics.sdstate.edu/idep93/).
