## Supplementary material for "Phenotypic and transcriptomic analysis reveals early stress responses in transgenic rice expressing Arabidopsis DREB1a": Table S1

| **Gene Name** | **Primer Name** | **Sequence (5'-3')** |
| --- | --- | --- |
| OsCAM1 | qOsCAM1-F | ACCAACCTCGGCGAGAA |
|  | qOsCAM1-R | CATGACCTTGACGAACTCC |
| OsWRKY76 | qOsWRKY76-F | ATCACGCTCGACCTCACCAAG |
|  | qOsWRKY76-R | AACTCCGGCGACGCGACCT |
| OsRbOHH | qOsRbOHH-F | GCAGGACTACTGGAAGTACGA |
|  | qOsRhOHH_R | GTCGTCGTAGATGTGGTTGAG |
| OsJAZ11 | qOsJAZ11-F | GCAGAGGCTATAATGAGGATGG |
|  | qOsJAZ11-R | CTTTGGCAAAATTGCCTAC |
| OsJAZ5 | qOsJAZ5-F | GTGTCCTTCTTCCATCTTCC |
|  | qOsJAZ5-R | ATTGGCCTGAGCAACAGGAA |
| OsFLS2 | qOsFLS2-F | AGTTCGCGTACATGAGGA |
|  | qOsFLS2-R | GGTGAAGAGCTCCATCG |
| OsbHLH48 | qOsbHLH148-F | ATTGCGGCTTGTGAAGTG |
|  | qOsbHLH148-R | ATCGCTGTCAGGCTGGTT |
| OsNAC59 | qOsNAC59-F | GGAAAACGACGGATCAGGAA |
|  | qOsNAC59-R | AGTACAGAAGTGCGGTCAAG |
| OsMAPKK17-18 | qOsMAPKK17-18F | AGATCGGAAGCTGTAATGGATG |
|  | qOsMAPKK17-18R | GGAGCTGTTCATCCACAAAGG |
| OsDERF5 | qOsDERF5-F | CAAGCTCAACTTCCCGG |
|  | qOsDERF5-R | TTCGAGCCGCAGTTCTC |
| OsDREB1a | qOsDREB1a-F | ATGTGCGGGATCAAGCAGG |
|  | qOsDREB1a-R | TCCACACCGTCTGGTGCT |
| 7Ubiquitin | q7Ubiq1445-F | TGGTCAGTAATCAGCCAGTTTG |
|  | q7Ubiq1520-R | CAAATACTTGACGAACAGAGGC |
