## Supplementary material for "Phenotypic and transcriptomic analysis reveals early stress responses in transgenic rice expressing Arabidopsis DREB1a": Table S2

| **Parameter** | **N_RT rep1** | **N_RT rep 2** | **N_CS rep1** | **N_CS rep2** | **T_RT rep1** | **T_RT rep2** | **T_CS rep1** | **T_CS rep2** |
| --- | --- | --- | --- | --- | --- | --- | --- | --- |
| **Globals** |  |  |  |  |  |  |  |  |
| Reference size | 375,049,285 | 375,049,285 | 375,049,285 | 375,049,285 | 375,049,285 | 375,049,285 | 375,049,285 | 375,049,285 |
| Number of reads | 48,505,594 | 38,443,370 | 46,438,394 | 39,719,254 | 43,101,920 | 47,086,070 | 42,139,082 | 38,515,704 |
| Average | 43,474,482 | | 43,078,824 | | 45,093,995 | | 40,327,393 | |
| Mapped paired reads | 46,779,108 (96.44%) | 37,005,302 (96.26%) | 44,717,370 (96.29%) | 38,413,656 (96.71%) | 41,447,690 (96.16%) | 46,355,280 (98.45%) | 40,923,840 (97.12%) | 36,763,238 (95.45%) |
| GC content | 53.06% | 52.98% | 52.69% | 53.02% | 53.54% | 53.62% | 52.55% | 52.24% |
| **Coverage** |  |  |  |  |  |  |  |  |
| Mean | 37.7874 | 30.1751 | 36.4708 | 31.2465 | 33.0932 | 36.978 | 33.6093 | 30.2478 |
| Standard deviation | 267.6963 | 212.5377 | 309.927 | 268.7895 | 280.9564 | 313.9022 | 288.124 | 253.8906 |
| **Mapping** |  |  |  |  |  |  |  |  |
| Mean Quality | 58.06 | 58.08 | 58.02 | 58.05 | 58.09 | 58.08 | 58.06 | 58.06 |

N_RT: non-transgenic at room tepretaure; N_CS: non-transgenic after cold-shock; T_RT: RD29a:DREB1a transgenic at room temperature; T_CS: RD29a:DREB1a transgenic after cold-shock
