## Supplementary material for "Phenotypic and transcriptomic analysis reveals early stress responses in transgenic rice expressing Arabidopsis DREB1a": Table S3

| **Gene construct in pNS64^1^** | **Gene** | **NCBI accession no.** | **Read Counts^2^** | | **Fold Change upon CS** |
| --- | --- | --- | --- | --- | --- |
|  |  |  | **T_CS** | **T_RT** |  |
| ZmUbi:NPTII | *NPTII* | KT184682.1 | 45,353 | 25,704 | 1.7 |
| 35Sppdk:GFP | *GFP* | U55762.1 | 42,362 | 34,775 | 1.2 |
| 35S:GUS | *GUS* | S94464 | 35,158 | 18,875 | 1.8 |
| RD29a:DREB1a | *DREB1A* | NM_118680.2 | 149 | 3 | 49.6 |
| GmHSP17.5E:pporRFP | *pporRFP* | DQ206380.1 | 6 | - | ~ |

^1^pNS64 harboring 4 gene constructs was used for Cre-*lox* mediated site-specific integration into rice genome. The resulting transgenic lines were used in this study

^2^Mapped read counts of cold-stressed (CS) or room temperature control (RT) transgenic lines (T)
